# Combining chromatography coupled SAXS and AI-predicted structures to dissect the mechanism of ParB1-*parS1* partition assembly formation

**DOI:** 10.1101/2024.06.25.600654

**Authors:** Anu Sodhi, Sneh Lata, Barnali N. Chaudhuri

## Abstract

Coupling of solution SAXS and AI-predicted structures can be a powerful strategy for delineating subtle conformational changes and self-association in protein switches. ParB, which is a condensate-forming DNA clamp that aids in bacterial chromosomal origin segregation, undergoes CTP-induced conformational switching to enable DNA sliding. The nature of *parS* DNA-induced conformational change in full-length ParB, and the structural features that govern self-association of ParB for partition assembly condensate formation, remains sparsely understood. We combined chromatography-coupled SAXS, rigid domains obtained from Alphafold model of ParB1 from *Vibrio cholerae*, and synthetic SAXS data describing known domain interfaces, to build integrative models of conformational states of full-length ParB1. These integrative models revealed how *parS1* DNA loading primed ParB1 for clamping and sliding. The CTPase domains in ParB1 were moved nearer upon DNA loading to facilitate clamping, and a lumen lined with a weak DNA binding site was formed below *parS1* binding site for capturing the sliding DNA. Furthermore, we showed that an N-terminal segment of ParB1 undergoes concentration-dependent oligomerization. An intrinsically disordered linker joining this oligomerization-prone N-terminal segment and the C-terminal domain curbs self-association of full-length ParB1, which is likely relevant for ParB1-mediated higher order partition assembly formation. To summarize, SAXS and Alphafold were effectively combined to provide unique insights into context-specific domain rearrangements and self-association in ParB1 for the mechanistic understanding of partition assembly formation.

## INTRODUCTION

Even though ATP or GTP-dependent protein switches are abundant in nature, only a handful of CTP switches are known (McLean and Le, 2023). Recently it was shown that ParB is a CTP switch that acts as a DNA clamp (Osorio-Valeriano et al., 2019; Soh et al., 2019; Osorio-Valeriano et al., 2021). ParB is a component of the bacterial chromosome segregation cassette that binds palindromic *parS* site(s) near the chromosomal origin (Funnell, 2016; Jalal and Le, 2020; Tišma et al., 2024; Livny et al., 2007). Multiple ParBs can spread around the *parS* site(s) to form a large partition assembly probably by sliding on the DNA and bridging (Funnell, 2016; Jalal and Le, 2020; Tišma et al., 2024; Sanchez et al., 2016; Walter et al., 2020; Balaguer et al., 2021; Antar et al., 2021; Guo et al., 2022; Tišma et al., 2022; Connolley et al., 2023; Tišma et al., 2023). Additionally, ParB participates in the loading of SMC proteins on DNA (Bock et al., 2022). Earlier, several mechanisms of DNA segregation have been proposed (Lim et al., 2014; Vecchiarelli et al., 2014). ParB forms CTP-dependent phase separated condensates (Guilhas et al., 2020; Babl et al., 2022). Biological roles of these condensates in ParB-mediated DNA segregation are just beginning to emerge (Azaldegui et al., 2021).

A canonical ParB is a dimer containing three domains (Funnell, 2016; Jalal and Le, 2020). The N-terminal domain of ParB/ParB-like protein binds and hydrolyze CTP (CTP-binding domain or CTPBD, Osorio-Valeriano et al., 2019; Soh et al., 2019; Osorio-Valeriano et al., 2021). A central helix-turn-helix motif-containing domain (HTHD) specifically binds palindromic *parS* DNA (Jalal et al., 2020). The C-terminal dimerization domain (CTDD) of ParB harbours a weak DNA-binding site (Fisher et al., 2017). Recent crystal structures of the nucleotide-bound and *parS*-bound truncated ParB family of proteins aided in understanding the CTP-dependent DNA clamping mechanism (Osorio-Valeriano et al., 2019; Soh et al., 2019; Osorio-Valeriano et al., 2021; Jalal et al., 2020; Jalal et al., 2021; Chu et al., 2024). Binding of two nucleotides at the dimer interface of CTPBD forms a locked “N-gate”, thus forming a closed ParB clamp (Osorio-Valeriano et al., 2019; Soh et al., 2019; Osorio-Valeriano et al., 2021; Jalal et al., 2021). This CTP-driven “N-gate” closure displaces *parS* DNA from the central HTHD, which is required for ParB sliding. CTP hydrolysis is catalysed by *parS*, suggesting a coupled conformational change in ParB (Jalal et al., 2020a). Structures of the full-length ParB in various conformations that are critical for understanding the domain rearrangements required for clamping-sliding have not been determined.

Artificial intelligence (AI) based structure prediction methods, such as Alphafold2, are now widely used, convenient tools for predicting 3-dimensional structures of folded proteins, including multi-domain and multimeric proteins, from primary sequences (Jumper et al., 2021; Mirdita et al., 2022; Agarwal and McShan, 2024). However, these predictions provide only one conformational state of the protein. Interconversions between conformational states associated with different stimuli are critical for functioning of protein switches (Alberstein et al., 2022). These context-dependent conformational states cannot be immediately deduced from one AI-predicted structure. Small angle X-ray scattering (SAXS) can be suitably combined with these AI-predicted structures for modelling various conformations associated with protein function (Brookes et al., 2023; Byer et al., 2023; Chinnam et al., 2023). Advanced methodologies, such as size exclusion column chromatography coupled SAXS (SEC-SAXS), and evolving factor analysis, can be particularly useful for separating scattering profile of the assembly component from SAXS data obtained from a mixture of components (Mathew et al., 2004; Meisburger et al., 2016). These methods can aid in better modelling of the conformational states of multi-component assemblies, such as a nucleoprotein complex. In addition, SAXS is a fitting tool for probing molecular features of proteins that can contribute to condensate formation by self-association (Martin et al., 2021).

We combined SEC-SAXS and AI-predicted structures to build integrative or hybrid models of the conformational states of ParB1 from *Vibrio cholerae* (vcParB1) and *parS1* (Fogel and Waldor, 2005; Yamaichi et al., 2007; Ramachandran et al., 2014; Niault et al., 2023). Comparison of these integrative models showed large conformational change in vcParB1 upon DNA loading, leading to repositioning of the CTPBD and formation of a lumen region below *parS1*-binding site for capturing the DNA. In addition, we show that a truncated vcParB1 containing the CTPBD and HTHD, and none of the putative intrinsically unstructured regions (IDR), oligomerize at high concentration. A flexible linker joining the CTDD and HTHD appears to restrain the self-associating tendency of full-length vcParB1. DNA-induced conformational changes and self-interaction of vcParB1 are discussed in the context of clamping-sliding and partition-assembly forming functions of vcParB1.

## RESULTS

### DNA binding induces conformational change in ParB1

Recently, it was shown that ParB families of proteins are CTP-dependent DNA clamps (Osorio-Valeriano et al., 2019; Soh et al., 2019; Osorio-Valeriano et al., 2021; Jalal et al., 2021). Partial mechanisms of conformational changes associated with DNA clamping were proposed based on the crystal structures of truncated ParBs (Osorio-Valeriano et al., 2019; Soh et al., 2019; Osorio-Valeriano et al., 2021; Jalal et al., 2020; Jalal et al., 2021; Chu et al., 2024). To understand the effect of DNA on the conformation of full-length vcParB1, we performed a series of SEC-SAXS runs for vcParB1, 16-meric palindromic *parS1* DNA, a non-specific DNA duplex of equal length (nsDNA16), and vcParB1-DNA assemblies (Figures 1A and S1). We experimentally confirmed that, like other reported ParBs, vcParB1 is a CTPase and can bind DNA (Figure S2). Based on the distribution of radii of gyration (R_g_) in the vcParB1 SEC-SAXS elution profile (Figure 1A), as well as singular value decomposition (SVD) surrounding the peak region (Figure S3A-B), an averaged SAXS profile (vcParB1-peak) was obtained. The vcParB1-peak profile was analysed to obtain size parameters (Table 1 and Figure S4). The eluted vcParB1 is a dimer (predicted mass: 59.5 kDa, theoretical monomeric mass: 34.1 kDa, Hajizadeh et al., 2018).

**Figure 1.**
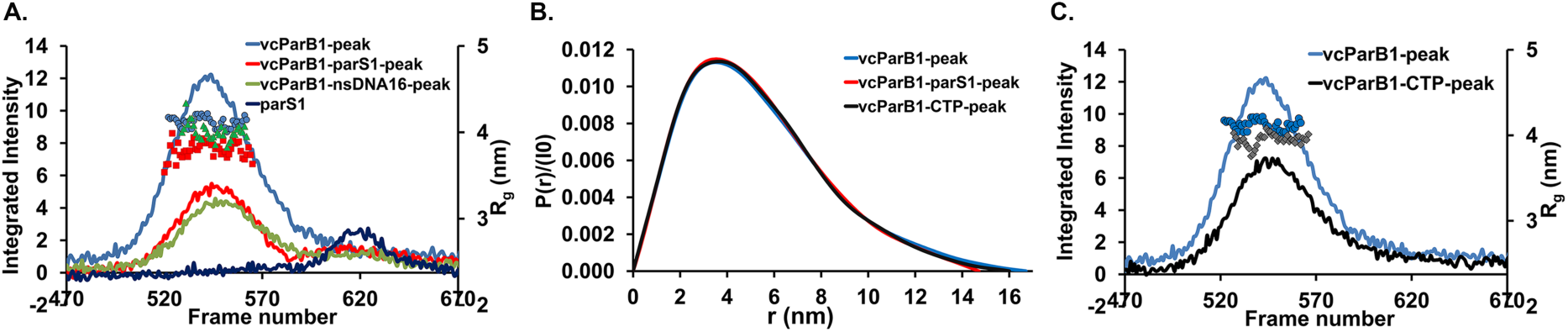
SEC-SAXS profiles of vcParB1 in unbound and bound states. (A) SEC-SAXS profiles, along with R_g_ distribution, for vcParB1 (solid blue line, R_g_ as blue circle), vcParB1-*parS1* (solid red line, R_g_ as red square) and vcParB1-*nsDNA16* (solid green line, R*_g_* as green triangle) are shown (x-axis: SEC-SAXS frame number; primary vertical axis (left): integrated intensity in arbitrary unit; secondary vertical axis (right): R_g_ in nm, calculated as a moving average of 5 frames). Elution profile for *parS1* (dark blue solid line) is shown. Elution profile for *nsDNA16*, which elutes at the same place as *parS1*, is not shown. (B) Normalized pair distribution functions (P(r)/I(0) *versus* r, P(r) is pair-distribution function, I(0) is forward scattering intensity and r is pairwise distance between atoms in nm) of vcParB1-peak (blue), vcParB1-*parS1*-peak (red) and vcParB1-CTP (black) are shown. (C) SEC-SAXS elution profiles of CTP-treated vcParB1 (solid black line, with R_g_ as grey diamond), and vcParB1 (solid blue line, with R_g_ shown as blue circle), are shown (plotted in the same way as in 1A).

**Table 1.**
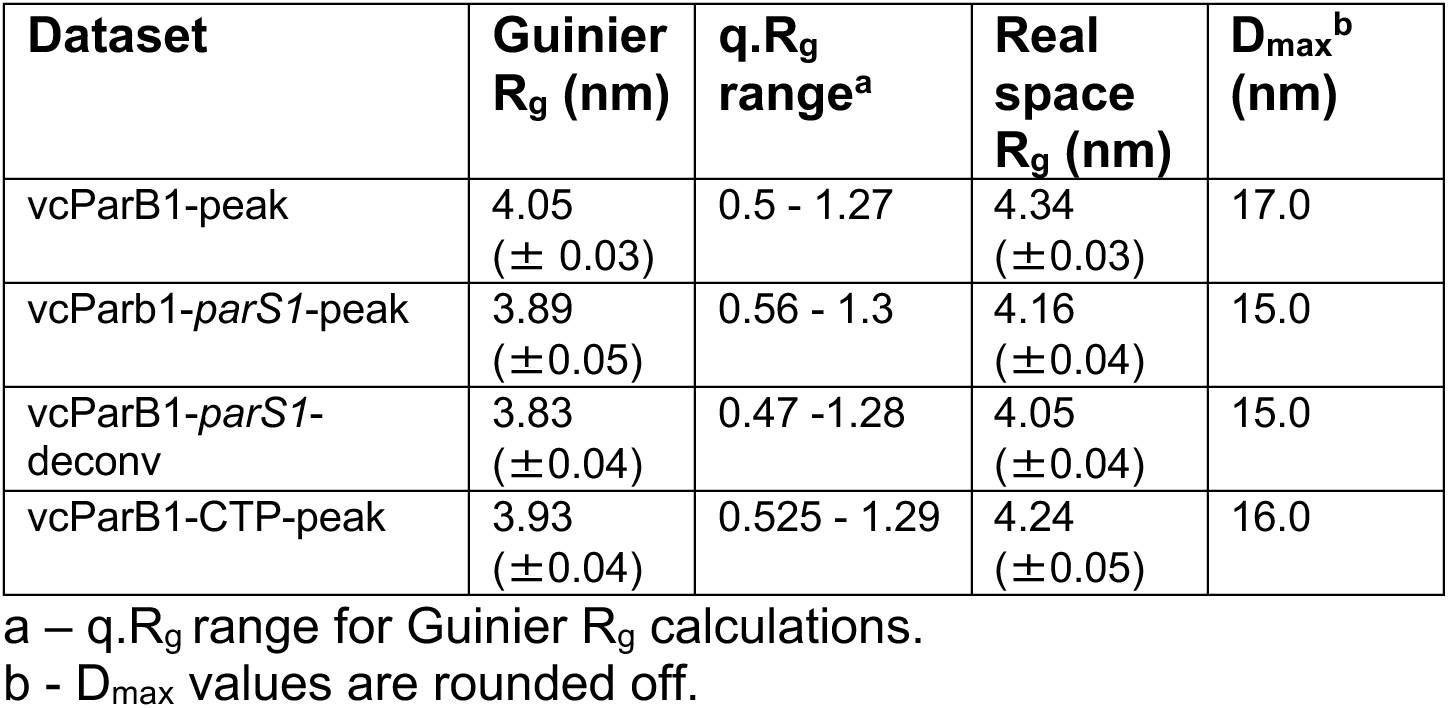
SEC-SAXS data analysis. BioXTAS RAW (Hopkins et al., 2024) and ATSAS (Manalastas-Cantos et al., 2021) were used for computing R_g_ and maximum particle diameter (D_max_).

| Dataset | Guinier $R_g$ (nm) | $q.R_g$ range<sup>a</sup> | Real space $R_g$ (nm) | $D_{max}$<sup>b</sup> (nm) |
| --- | --- | --- | --- | --- |
| vcParB1-peak | 4.05<br>($\pm 0.03$) | 0.5 - 1.27 | 4.34<br>($\pm 0.03$) | 17.0 |
| vcParb1-*parS1*-peak | 3.89<br>($\pm 0.05$) | 0.56 - 1.3 | 4.16<br>($\pm 0.04$) | 15.0 |
| vcParB1-*parS1*-deconv | 3.83<br>($\pm 0.04$) | 0.47 - 1.28 | 4.05<br>($\pm 0.04$) | 15.0 |
| vcParB1-CTP-peak | 3.93<br>($\pm 0.04$) | 0.525 - 1.29 | 4.24<br>($\pm 0.05$) | 16.0 |

a – $q.R_g$ range for Guinier $R_g$ calculations.
b - $D_{max}$ values are rounded off.

A comparison of the SEC-SAXS profiles of vcParB1, vcParB1-*parS1*, vcParB1-*nsDNA16*, *parS1* and *nsDNA16*, showed that the eluted DNA peaks were separated from the eluted protein and protein-DNA complex peaks (Figure 1A). Overall R_g_ distribution surrounding the peak region showed that the average size of the vcParB1-*parS1* complex, and that of the vcParB1-nsDNA16 complex to a lesser extent, is reduced when compared to the average size of vcParB1 (Figure 1A). A similar *parS*-induced shrinking in mycobacterial ParB was reported earlier (Chaudhuri and Dean, 2011). This shift in R_g_ is an indication of conformational change in vcParB1 upon vcParB1-*parS1* assembling.

To obtain SAXS profile of the vcParB1-*parS1* complex from the SEC-SAXS data collected from a mixture of protein, DNA and DNA-protein complex, we performed REGularized Alternating Least Squares (REGALS) decomposition (Meisburger et al., 2021). REGALS of vcParB1-*parS1* elution profile provided 3 components (Figure S5A-C). The major component (vcParB1-*parS1*-deconv), which was most compact in the Kratky plot, was assigned to be the protein-DNA complex (Figure S5A). SVD analysis of the vcParB1-*parS1* peak region identified a segment containing one component, which was averaged (vcParB1-*parS1*-peak; Table 1 and Figure S3C-D), and used for subsequent analysis along with the vcParB1-*parS1*-deconv profile (Table 1). A comparison of pair distribution functions (PDF) obtained from the vcParB1-*parS1*-peak/vcParB1-*parS1*-deconv and vcParB1 profiles showed that the vcParB1-*parS1* is ∼ 20 Å smaller in size than vcParB1, further supporting conformational change in vcParB1 upon *parS1* binding (Table 1 and Figure 1B).

To find the effect of CTP on vcParB1 conformation, we performed SEC-SAXS experiments with CTP-treated vcParB1 in the presence of mobile buffer supplemented with 1.2 mM of CTP. SVD analysis aided in obtaining averaged intensity profile of nucleotide-treated vcParB1 (vcParB1-CTP-peak) from a set of adjacent frames (Figure S3E-F). A small reduction in the size of vcParB1 upon CTP binding suggested that, like *parS1*, CTP induces conformational change in vcParB1 (Table 1 and Figure 1B-C).

### Structural modeling of vcParB1 conformational states

Even though the crystal structures of truncated domains of ParB/ParB-like proteins, and their complexes, yielded valuable insights into the ParB DNA clamping mechanism (Osorio-Valeriano et al., 2019; Soh et al., 2019; Osorio-Valeriano et al., 2021; Jalal et al., 2020; Jalal et al., 2021), structures of full-length ParB, and ParB-*parS* complex, are required to fully comprehend the mechanism of clamping. Probably due to the presence of IDRs, crystal structures of full-length ParBs are unavailable. Cryo-EM can provide structures of full-length ParB in different conformational states, which may not represent a physiologically relevant solution state (Bock et al., 2022a). SAXS can provide information about low-resolution solution shapes of proteins in different biologically relevant conditions that can be combined with AI-predicted structures to model condition-dependent conformational states.

We used Alphafold multimer in colabfold (Mirdita et al., 2022), with custom templates, to predict structures of dimeric vcParB1 (Figure S6A). Poorly defined segments with low predicted local distance difference test (pLDDT) scores in these structures, one in the N-terminal region and another short linker joining CTPBD and HTHD, appear to correlate well with predicted IDRs (Figure S6A-B). Structures of CTPBD and HTHD of vcParB1 were comparable to the template structures (Figure S6C-D). Structure of C-terminal CTDD of vcParB1 was strikingly similar to the NMR structures of CTDD of ParB from *B. subtilis* (Fisher et al., 2017), with which it bears low sequence identity (Figure S6E). SAXS profile calculated using this predicted model of vcParB1, which is in “N-gate” closed form, does not match any of our experimental SAXS profiles (Figure S6F-H). The AI-predicted structure must be altered to obtain conformations of vcParB1 that are consistent with SAXS data collected under different conditions.

To obtain the average solution structures of vcParB1 and vcParB1-*parS1* assembly, AI-predicted structures of rigid domains of vcParB1 (and *parS1* for the nucleoprotein assembly) were combined with experimental SAXS data, synthetic SAXS data based on known domain interfaces, 2-fold symmetry constraint, and additional distance restraints, for structural modeling (see methods section, Petoukhov et al., 2012). Consistent hybrid structural models corresponding to vcParB1-peak, vcParB1-*parS1*-peak and vcParb1-parS1-deconv datasets were obtained with good fit to SAXS profiles (Figures 2A-E, S7 and S8). 20 V-shaped hybrid structures of vcParB1 were clustered into two conformational groups by manual inspection that cannot be distinguished by SAXS (cluster I and cluster II; 10 structures in each cluster; Figure 2B-C). We concluded that unbound vcParB1 adopted open V-shaped forms in solution (Figure 2A-C).

**Figure 2.**
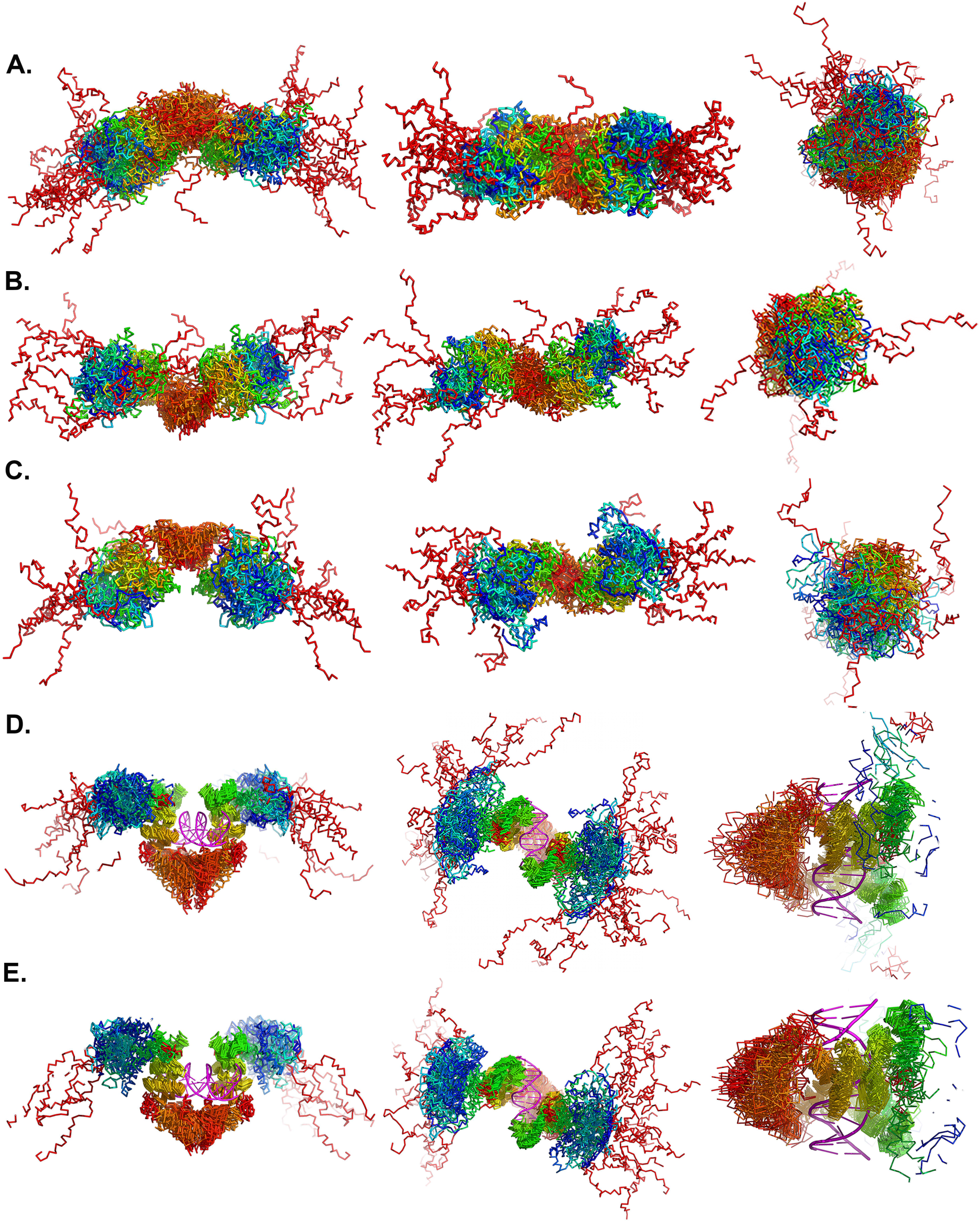
Hybrid structural models of vcParB1 and vcParB1-*parS1* assembly. Three different views of (A) 20 superposed hybrid structures of vcParB1 (vcParB1-peak data), (B) 10 superposed hybrid structures of vcParB1 cluster I, (C) 10 superposed hybrid structures of vcParB1 cluster II, (D) 20 superposed hybrid structures of vcParB1-*parS1* (vcParB1-*parS1*-peak data), (E) 20 superposed hybrid structures of vcParB1-*parS1* (vcParB1-*parS1*-deconv data). All structures are shown in ribbon (in rainbow, the N-terminal IDR is shown in dark-red). Centrally located CTDD was used for all alignments in A-C. The structure of *parS1* was kept fixed during the modelling of vcParB1-*parS1* complexes, which are shown without alignment in figure D-E.

The *parS1*-bound hybrid models of vcParB1 revealed a lumen region surrounded by the HTHD, linker region and CTDD (Figure 2D-E). The CTPBD in these models are not in “N-gate closed” form or as proximal as observed in the structure of cvParB1trunc-*parS* (pdb code: 6t1f). An attempt to model vcParB1-*parS1* complex with distance restraints to bring the CTPBD closer led to high χ^2^ values (Figure S7D).

Despite good χ^2^ values (Figure S7E), much variability was observed in the hybrid structures of vcParB1 modeled against vcParB1-CTP-peak data. Therefore these models are not discussed. When “N gate closed” condition was imposed as an additional synthetic data during modeling, hybrid structures of vcParB1 yielded high χ^2^ values (Figure S7F). Therefore, we concluded that the CTP-treated vcParB1 is in an “N-gate” open average conformation in solution.

### A truncated form of vcParB1 devoid of the intrinsically unstructured regions is highly heterogeneous

Surprisingly, the SEC-SAXS elution profile of a truncated construct of vcParB1 (vcParB1trunc) without the IDR and CTDD showed noticeable heterogeneity in size (Figure 3A). Distribution of R_g_ around the eluted peak with a distinct slope, and SVD analysis, indicated the presence of multiple components that are being separated while eluting (Figure 3A-B) (Martin et al., 2021). As it was not possible to obtain an averaged SEC-SAXS profile of vcParB1trunc due to a large variation in R_g_ across the peak, a number of SAXS profiles averaged over a few frames from different sections of the eluted peak were used for further analysis (Figure S9A). Kratky plots obtained from these SAXS profiles (sequentially numbered from I to VI from left to right) suggested presence of folded proteins (Figure 3C). Comparison of PDFs obtained from these SAXS profiles showed a large variation in D_max_, further supporting heterogeneity in vcParB1trunc (Figure 3D).

**Figure 3.**
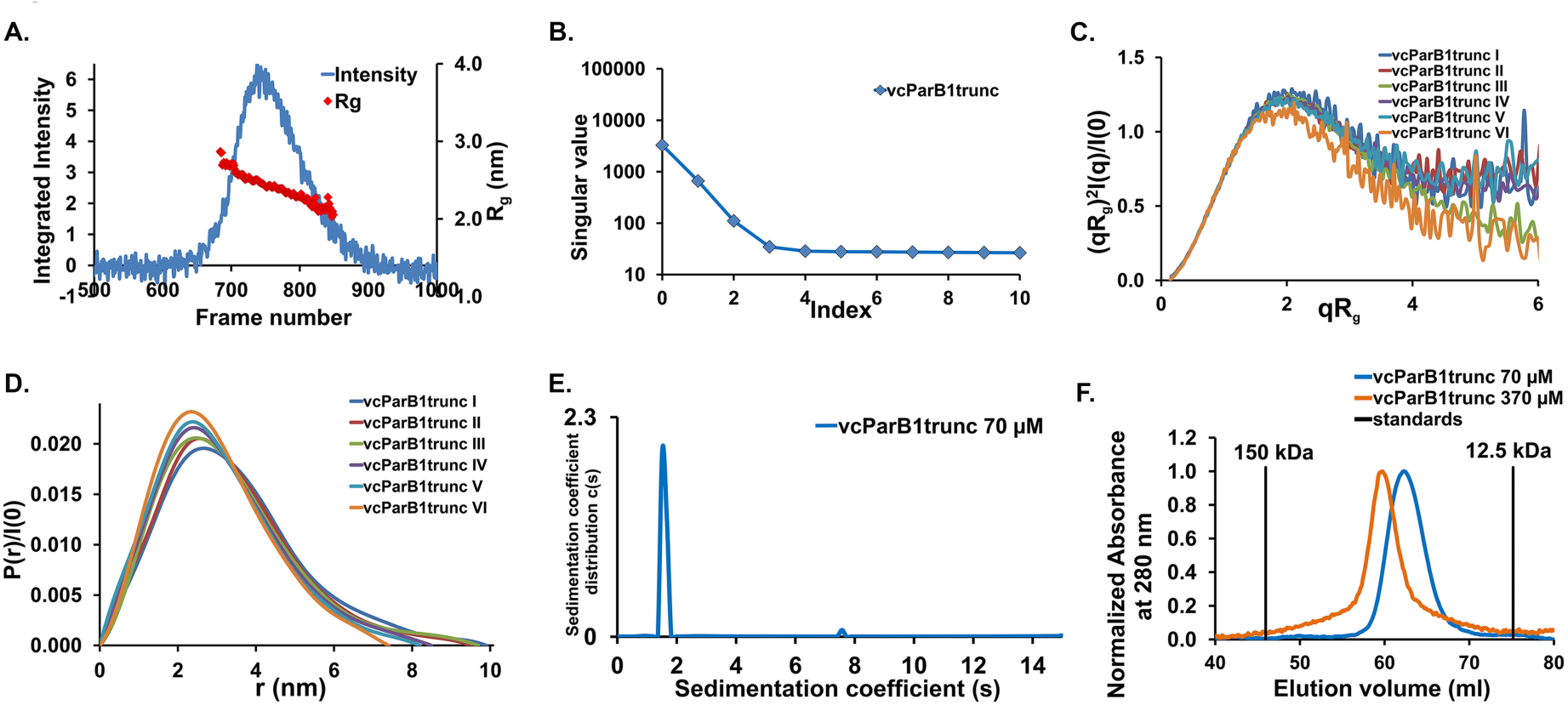
Purified vcParB1trunc is heterogeneous in nature. (A) SEC-SAXS profile of vcParB1trunc with integrated intensity (primary vertical axis, blue line, arbitrary unit) and moving average R_g_ over 5 frames (secondary vertical axis, red diamond, in nm) *versus* frame numbers. (B) Result of SVD of the vcParB1trunc SEC-SAXS dataset encompassing the peak region. Semilog plot of singular value *versus* index is shown. (C) Dimensionless Kratky plots, (qR_g_)^2^I(q)/I(0) *versus* qR_g_ (I(q) is intensity in arbitrary unit and q is momentum transfer), and (D) normalized PDF (normalized by I(0)) of averaged SAXS profiles of segments obtained from the above dataset, with segment ranges I to VI (early to late eluting segments). PDFs were calculated and shown as described in figure 1B. (E) Plot of sedimentation coefficient distribution c(s) *versus* sedimentation coefficient (s) is shown for vcParB1trunc (blue). (F) SEC profiles (normalized to maximum value of 1) of vcParB1trunc from the same batch at two concentrations are shown in blue (70 μM) and orange (370 μM). Vertical black lines show the peak positions of cytochrome c (12.5 kDa) and alcohol dehydrogenase (150 kDa).

This trimmed vcParB1trunc showed weak CTPase activity and DNA binding abilities (Figures S2C-D and S9B). Analytical ultracentrifugation (AUC) confirmed that purified vcParB1trunc (∼ 70 μM) is a monomer (estimated mass 23.4 kDa, theoretical mass 23.9 kDa; Figure 3E). A repeat of this AUC run in the presence of 1 mM CTP rendered the data not analyzable, which is likely due to CTP-induced aggregation. A high concentration (∼ 386 μM) of vcParB1trunc was used for the SEC-SAXS experiment, which is unsuitable for AUC. Therefore, a size-exclusion column chromatography (SEC) run with vcParB1trunc at two concentrations (∼ 70 and ∼ 370 μM) was performed, that was consistent with oligomerization of vcParB1trunc at higher concentration (Figure 3F). According to previous reports, CTPBD of ParBs can oligomerize in many ways, such as a dimer and a tetramer (Soh et al., 2019; Chen et al., 2015). At high concentration, vcParB1trunc probably exists as a mixture of oligomers that are formed by intermolecular interactions.

### Elevated flexibility in the linker region promotes self-association of vcParB1

Next, we asked, what molecular features could be responsible for balancing between the intra-molecular clamp formation within the dimeric vcParB1 and intermolecular self-association that is required for higher order partition assembly formation? A short flexible linker region has been identified in many ParB homologs that join the CTDD and HTHD (Figures S6B and S10A). A number of predicted ParB structures downloaded from the Alphafold2 database (alphafold.ebi.ac.uk) showed very low pLDDT score in this linker region, further supporting predicted flexibility in this region (Figures 4A and S6A). We hypothesized that this linker is restricting the N-terminal region of wild-type vcParB1 dimer from self-interaction with neighboring dimers by limiting the rotational freedom around the linker. To test this hypothesis, we designed a glycine-rich triple mutant (vcParB1-linker; P235G, D237G, K239A, Figure S10A) for elevating flexibility in this linker region. We expect that a highly flexible linker will allow the N-terminal segment of vcParB1-linker to liberally rotate about the linker to sample wider conformational space, thus enhancing chances of intermolecular interactions for vcParB1-linker when compared to the wild-type.

**Figure 4.**
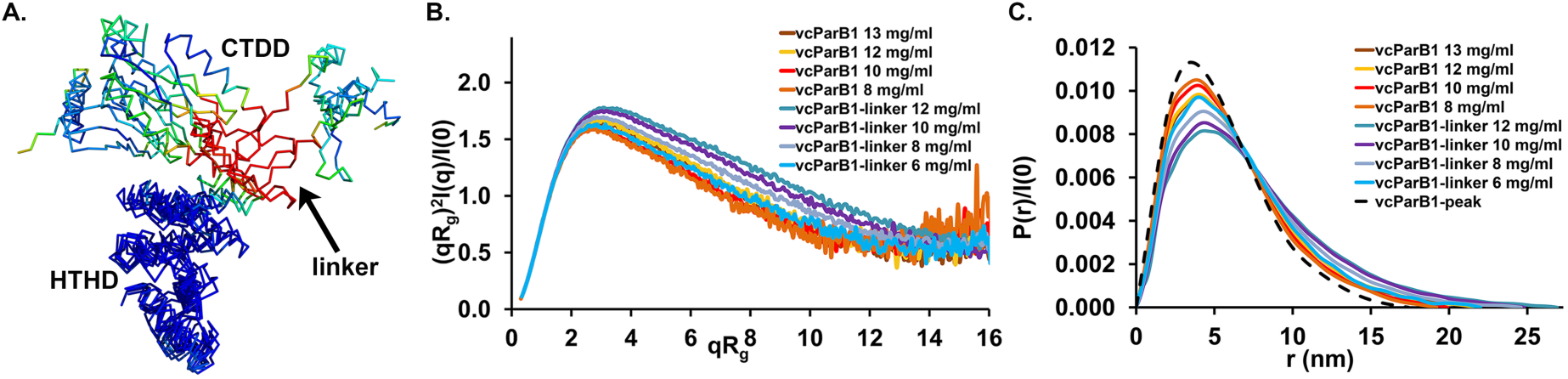
Triple-mutant vcParB1-linker is more association-prone than vcParB1. (A) Superposed Alphafold predicted structures of vcParB1 and ParB homologs (Uniprot accession numbers: Q9PB63, Q9RYD8, Q87BY1, Q92JI0, Q1CVJ4, Q9JW77), coloured by pLDDT score (90 in blue to 50 in red), are shown. Predicted structures were superimposed using the Cα atoms of HTHD. The CTPBD domains are not shown. (B) Dimensionless Kratky plots (as described in Fig. 3C) and (C) normalized PDFs (as described in Fig. 1B) of SAXS data obtained from vcParB1 and vcParB1-linker. Real space R_g_ and I(0) values from the PDFs were used for calculating these Kratky plots. PDF obtained from the vcParB1-peak SEC-SAXS data is plotted as a dashed black line in 4E.

Purified vcParB1-linker showed weak CTPase activity (Figure S10B). Cumulative polydispersity indices obtained from dynamic light scattering experiments supported higher level of intermolecular interactions in vcParB1-linker compared to vcParB1 (Figure S10C).

To further compare self-associating properties of vcParB1 and vcParB1-linker, we obtained batch-mode SAXS data. Kratky plots obtained from these SAXS data suggested high level of self-association (as opposed to self-avoiding behavior, Martin et al., 2021) in the vcParB1 constructs (Figure 4B). D_max_ obtained from the vcParB1-linker datasets were consistently higher than that from the vcParB1 datasets, suggesting higher degree of self-association in the linker mutant (Figure 4C). To quantitate self-interaction in vcParB1 and vcParB1-linker, we calculated second virial coefficient (B22) from SAXS data (Pabit et al., 2009). B22 for vcParB1 and vcParB1-linker were estimated to be - 3.99 X 10^-4^ mol.mL.g^-2^ and - 4.66 X 10^-4^ mol.mL.g^-2^, respectively. Negative B22 values imply attractive interaction, and it appears from the B22 values that the vcParB1-linker is more association-prone than vcParB1 within our experimental concentration regime. Thus, imparting more flexibility in the linker region made vcParB1 more association-prone.

## DISCUSSION

Since the recent serendipitous discovery that ParB is a CTP switch and a DNA clamp, number of CTP switches reported in the literature is steadily growing (McLean and Le, 2023). However, a lack of structural data on the key conformations of full-length ParB in various biologically relevant conditions prevented thorough mechanistic understanding of the clamping-sliding process, which is addressed here.

SEC-SAXS revealed that conformational state of the vcParB1 changed in the presence of cognate *parS1*, and in the presence of nucleotide (Figure 1A, 1C). Next, rigid domains obtained from AI-predicted structures of vcParB1 were combined with average SEC-SAXS profiles, and with synthetic SAXS data based on known domain interfaces, to model solution conformational states of vcParB1 and vcParB1-*parS1* (Figure 2). SVD and REGALS tools were used for vetting the SEC-SAXS profiles that were used for structural modeling (Figures S3 and S5). We tested alternative hypotheses about domain orientations by using relevant synthetic data and/or distance restraints during modeling (Figure S7). Even though such structural models built using sparse data are not suitable for fine structural analysis, they can be powerful reporters of domain movements and interface formation in solution (Byer et al., 2023; Chinnam et al., 2023).

While consistent structural models of vcParB1-CTP were not obtained, our results agree with the expectation that considering the cytosolic concentration of CTP in bacteria (∼ 0.3 mM, Buckstein et al., 2008), majority of intracellular ParB will be in a CTP-bound, “N-gate” open form (Osorio-Valeriano et al., 2019). Average solution structures of vcParB1 in DNA-bound and DNA-free forms delineated a probable pathway for *parS1*-induced changes in vcParB1 that can lead to subsequent clamping and sliding (Figure 5A). In our structural models, the two CTPBD domains where moved nearer upon ParB1-*parS1* assembling to facilitate clamp closure, and a lumen poised for capturing the dislodged *parS1* was formed in ParB1 beneath the bound DNA, thus priming the ParB1 clamp for sliding (Figure 5A-C).

**Figure 5.**
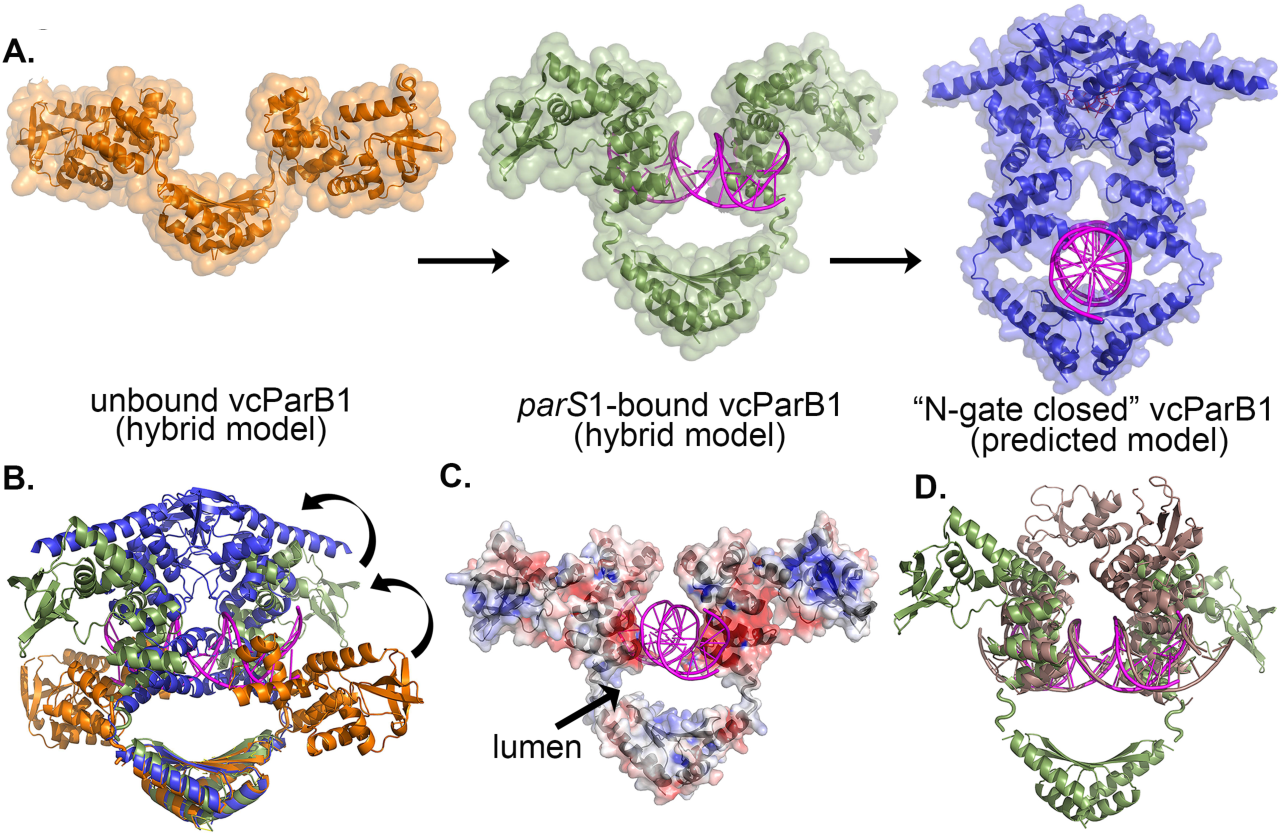
Conformational states of vcParB1. (A) Representative hybrid models of vcParB1 dimer (cluster I) in orange and vcParB1-*parS1* complex in green, with *parS1* in magenta, are shown as cartoons. AI-predicted structure of vcParB1 (blue), is shown as a cartoon. Two docked nucleotides in the CTPBD (red sticks), and a *parS1* DNA docked in the lumen region (magenta), are shown with the predicted structure of vcParB1 (blue). The *parS1* DNA is tightly fitted in the lumen, with steric conflicts. Further conformational changes in the lumen region will be required for accommodating the sliding DNA. Semi-transparent surfaces are shown along with all the cartoons. (B) Superposed representative structures of vcParB1 (orange), vcParB1-*parS1* (green, *parS1* in magenta) and AI-predicted model of vcParB1 (blue) are shown. The CTDD was used for structural alignment. Black curly arrows show the movement of CTPBD between different conformers of vcParB1. (C) Electrostatic potential mapped on the surface of vcParB1-*parS1* (shown as semi-transparent surface, along with the cartoon of the structure in grey, *parS1* in magenta). (D) Cartoons of a hybrid structure of vcParB1-*parS1* (green, *parS1* in magenta,) superposed on the crystal structure of cvParBtrunc-*parS* (pink, superposed using the HTHD; pdb code 6t1f). The flexible N-terminal region is not included in the cartoons for clarity.

Unlike in our hybrid models, the CTPBD domains in the crystal structure of truncated cvParB-*parS* are in physical proximity (Figure 5D). Crystallization probably favored a compact form of ParB-*parS*. These conformations could be lowly populated, and therefore not captured by our average solution structure modeling approach. The CTPBD in our hybrid model of vcParB1-*parS1* dimer is available for either intramolecular (required for clamping using a specific interface) or intermolecular (higher order self-association for partition assembly formation) interactions.

How self-association of different segments of ParB are modulated for partition assembly formation is an intriguing question. We used SAXS, which is an apt technique for understanding self-assembling in condensate-forming proteins (Martin et al., 2021), to probe and quantitate self-association in vcParB1. We note that SAXS is a weight-average method and presence of a small amount of higher order oligomer can change the SAXS profile significantly. A trimmed vcParB1trunc, which is folded and functional (Figures 3C, S2C-D and S9B), can oligomerize in a concentration-dependent manner. Mutational analysis suggests that the linker region joining the HTHD and CTDD is limiting the higher order self-assembling propensity of full-length vcParB1 (Figure 4). Linker conformations probably pose physical barrier to rampant self-association of vcParB1 dimer with other dimers by making the interaction interfaces of the N-terminal segments less accessible for intermolecular interactions.

Like other ParBs (Guilhas et al., 2020; Babl et al., 2022), vcParB1 form condensates (unpublished results). Therefore, self-associating propensity of vcParB1 is hardly surprising. Oligomerization of domains, amongst other factors, is a known influencer of condensate formation in multi-domain proteins (Mohanty et al., 2022). Since local concentration of ParBs can be as high as 500 μM within the partition assembly (Lim et al., 2014), observed oligomerization of vcParB1trunc at high concentration might be relevant for partition assembly condensate formation. The linker region likely helps to keep full-length vcParB1 in open state in the cytosol. On the other hand, within the highly crowded partition assembly, self-association of vcParB1 domains might prevail to aid in the formation and maintenance of the partition assembly. It appears that CTP is a driver of ParB condensate formation (Babl et al., 2022; Balaguer et al., 2021; Zhao et al., 2024). How the clamping-sliding and self-association of ParBs are combined for partition assembly formation, and how CTP supports these processes, are key questions remains to be answered.

## Conclusion

Contemporary protein structure prediction methods, such as Alphafold, can provide high quality structural models of proteins, including protein switches. However, the predicted single conformer Alphafold models are inadequate for mechanistic understanding of protein switching function in terms of their conformational changes in solution. We combined Alphafold model with SAXS data collected under different conditions to build conformational states of ParB1 DNA clamp. These conformational states showed how DNA loading prepares ParB1 for DNA-clamping and subsequent sliding. Further, biophysical studies on mutated ParB1 constructs suggested how ParB1 could intrinsically modulate self-association for partition assembly formation. The presented approach can be generalized for understanding molecular mechanisms of protein conformational switching and self-association.

## Materials and Methods

### Protein expression and purification

Either pET15b or pET28a vectors containing genes of target proteins (supplementary material **S1**) were transformed into BL21(DE3) competent cells. Cells were grown at 37° C in Luria-Bertani broth (Difco, Fisher scientific, cat 244620) with added antibiotics (100 µg/ml of ampicillin, Sisco Research Laboratory, cat 61314 or 50 µg/ml kanamycin monosulfate, Gold Biotechnology, cat K-120-5) till OD_600_ reached 0.6. Next, Isopropyl-β-D-thiogalactoside (IPTG, 1 mM; Gold Biotechnology, cat I2481C50) was used for induction and cells were grown for another 15 hours at 25° C. Cells were harvested and stored at - 80° C.

For protein purification, cells were lysed by sonication (Sonics Vibra-Cell, VCX750, Sonics & Material) in the lysis buffer containing 100 mM Tris.HCl (Sigma-Aldrich cat T6066) pH 7.5, 1 M NaCl (Sigma-Aldrich cat S3014), 5 % glycerol (Sigma-Aldrich cat G7757), 5 mM β-mercaptoethanol (BME, Sigma-Aldrich cat M6250), 0.2 mM ethylenediaminetetraacetic acid (EDTA, Sigma-Aldrich cat EDS) and protease inhibitor cocktail (Sigma-Aldrich, cat P8849). Proteins were purified using immobilized metal affinity column chromatography (IMAC sepharose 6 fast flow, Cytiva, cat 17092107), followed by ion exchange column chromatography (Q sepharose Fast Flow, Cytiva, cat 17051010) and size-exclusion column chromatography (HiLoad 16/600 Superdex 200 pg, Cytiva, cat 28989335). Proteins were loaded to Ni^+2^-charged IMAC sepharose fast flow column in lysis buffer. Unbound and weakly bound proteins were eluted by wash buffer (20 mM Tris.HCl pH 7.5, 1 M NaCl, 5 % glycerol), followed by a binding buffer containing 20 mM Tris.HCl pH 7.5, 300 mM NaCl, 5 % glycerol and 20 mM imidazole (Sigma-Aldrich cat I2399). Bound proteins were eluted in the binding buffer with increased concentration of imidazole in steps up to a maximum concentration of 500 mM. Next, the collected protein fractions were pulled and dialysed in the ion exchange buffer (20 mM Tris.HCl pH 7.5, 20 mM NaCl, 5 mM BME, 5 % glycerol, 1 mM EDTA). A gradient of 20 mM to 500 mM NaCl was used for eluting the protein. Buffer for the size-exclusion column chromatography run contained 20 mM Tris.HCl pH 7.5, 100 mM NaCl, 5% glycerol, 5 mM BME and 1 mM EDTA. Purity levels of the proteins were evaluated by sodium dodecyl sulfate-polyacrylamide gel electrophoresis (SDS-PAGE). Fractions from the peak region were collected and further concentrated. Trypsin digested (in-gel digestion, Trypsin singles, proteomics grade, cat T7575) proteins were further confirmed by MS/MS ion search from one or more peptides in MASCOT database server (Perkins et al., 1999). Protein concentrations were determined using BCA assay (Pierce BCA protein assay kit, cat 23225).

### Oligonucleotide preparation

Commercially synthesized oligonucleotides were obtained in PAGE-purified form from Sigma-Aldrich oligonucleotide synthesis facility (supplementary material **S1**). Equimolar concentrations of oligonucleotides were mixed and annealed by heating at 95°C (10 minutes), followed by cooling at room temperature (45 minutes) and at 4°C (overnight).

### CTPase activity assay

To detect CTPase activity of vcParB1, malachite green assay (Malachite green oxalate salt, Sigma-Aldrich, cat M6880, Ammonium molybdate tetrahydrate, Sigma, cat 09878) was performed. Standard curves were prepared using potassium phosphate monobasic (Sigma Aldrich, cat P5379). Protein and cytidine triphosphate (CTP; Thermo scientific, cat R0451, 1 mM) were incubated in the reaction buffer (20 mM Tris.HCl pH 7.5, 150 mM NaCl, 5 mM MgCl_2_) for 60 minutes at room temperature. OD_650_ was measured in a plate reader (Thermo Fisher scientific, Multiskan Go) after adding the malachite green working reagent and sodium citrate. OD obtained from the reaction buffer with only CTP was used as blank for subtraction. Next, these experiments were repeated in the presence of oligonucleotides (*parS1* or *nsDNA16*, supplementary material **S1**). All experiments were performed three to four times and plotted in Excel (Microsoft).

### Biolayer interferometry

ParB1-DNA interactions were detected by biolayer interferometry in our in-house Octet K2 (ForteBio, USA). Biotinylated duplex DNA (bt-*parS1*, Supplementary material **S1**) containing a single *parS1* sequence, and non-specific DNA (bt-*nsDNA*), were used for coating streptavidin biosensor (SA biosensor, Sartorius). Protein concentrations were optimized before the final runs in the reaction buffer containing 50 mM Tris.HCl pH 8.0, 150 mM NaCl, 5 mM MgCl_2_, 1 mM BME, 0.05 % Tween20 (Sigma-Aldrich, cat P9416), 0.5 % Bovine Serum Albumin (Sigma-Aldrich, cat A2153). All experiments were done in triplicate. Data analysis was performed using the software Data Analysis HT 11.1 (Sartorius). A heterogeneous binding model was used for data analysis that provided two on-rates and two off-rates. It was not possible to obtain reliable dissociation constant due to much variability in the data, which is probably due to the self-associating nature of vcParB1.

### Analytical ultracentrifugation

Analytical ultracentrifugation (AUC) runs were performed in duplicate at 42000 rpm at 20° C using ProteomeLab XL-I protein characterization system (Beckman coulter). An AN-50 Ti rotor was used and scans were taken at 280 nm. Purified protein was used for the AUC experiments with the following reference buffer: 20 mM Tris.HCl pH 7.5, 100 mM NaCl, 1 mM EDTA, 5 % glycerol, 5 mM BME. SEDFIT was used for the data analysis (Schuck, 2000).

### Dynamic light scattering

Dynamic light scattering (DLS) experiment was performed using Prometheus Panta (NanoTemper technologies), with laser wavelength 405 nm, available for demonstration. Purified proteins were used in a buffer containing 20 mM Tris.HCl pH 7.5, 100 mM NaCl, 1 mM EDTA, 5% glycerol, 5 mM BME at 20° C temperature for the DLS runs. Data analysis was performed using the Panta analysis software available with the instrument (NanoTemper technologies).

### Analytical size exclusion column chromatography

Size exclusion column chromatography runs were performed using HiLoad 16/600 Superdex 75 pg column (Cytiva) at 0.3 ml/min flow rate and at 10° C temperature. The following buffer was used for the SEC runs: 20 mM Tris.HCl pH 7.5, 100 mM NaCl, 1 mM EDTA, 5% glycerol, 5 mM BME. Prior to the experiment, the column was standardized using the gel filtration marker kit MWGF200 (Sigma-Aldrich). Excel (Microsoft) was used for plotting the results.

### SAXS and SEC-SAXS experiments

Purified proteins and protein-DNA complexes were snap-frozen under liquid nitrogen and shipped to the synchrotron site in a dry shipper. All data were collected at the beamline BM29, European Synchrotron radiation facility (ESRF), Grenoble, France (Tully et al., 2023). Experimental parameters are presented in **Table 2**.

**Table 2.** Data collection parameters for SAXS and SEC-SAXS experiments.

| Parameters | SAXS |
| --- | --- |
| Experimental station | BM29, ESRF |
| Wavelength ( Å ) | 0.999 |
| Detector distance (m) | 2.81 |
| Exposure time (sec) | 1 *per* frame (10 frames) |
| Protein buffer | 20 mM Tris pH 7.5, 100 mM NaCl, 5% glycerol, 1 mM EDTA, 5mM BME, 5mM MgCl<sub>2</sub> |
|  | **SEC-SAXS** |
| Experimental station | BM29, ESRF |
| Wavelength ( Å ) | 0.999 |
| Detector distance (m) | 2.81 |
| Exposure time (sec) | 2 *per* frame, ~1000 frames |
| SEC SAXS setup | AKTA Ettan system (GE healthcare) |
| SEC column | Superdex200 increase 3.2/300 |
| Running buffer | 20 mM Tris pH 7.5, 100 mM NaCl, 2 % or 5% glycerol, 1 mM EDTA, 5 mM BME or 2 mM DTT, 5 mM MgCl<sub>2</sub> (1.2 mM CTP added in buffer for the experiments with CTP-treated protein) |

### SAXS and SEC-SAXS data analysis

Resultant SAXS and SEC-SAXS data were analysed using BioXTAS RAW 2.2.1 (Hopkins, 2024) and ATSAS suite of software (Manalastas-Cantos et al., 2021). Following buffer subtraction, peak regions of protein and protein-DNA complexes from the SEC-SAXS elution profiles were selected based on SVD analysis using BIOXTAS RAW (Hopkins, 2024). Averaged scattering profiles were used for subsequent analysis. Regularized alternating least squares (REGALS, Meisburger et al., 2021) analysis was performed for the SEC-SAXS dataset of the vcParB1-*parS1* complex. Pair distribution functions were calculated using GNOM (Manalastas-Cantos et al., 2021) Second virial coefficients (B22) were estimated from concentration series of batch mode SAXS data (Pabit et al., 2009), using the following equation –

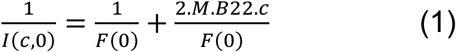

I(c,0) is the forward scattering intensity at concentration c, M is mass and F(0) is form factor at zero scattering angle. CRYSOL was used for calculating theoretical scattering profiles from atomic coordinates (Manalastas-Cantos et al., 2021). Excel (Microsoft) was used for preparing the graphs and analysing data.

### AI-based structure prediction

Structural models of full-length, dimeric vcParB1 were obtained from Colabfold (Mirdita et al., 2022), using Alphafold Multimer v3 option with custom templates. Known crystal structures of C-terminally truncated ParB from *Caulobacter vibrioides* (cvParBtrunc, ∼ 39 % sequence identity with vcParB1 for residue coverage of ∼ 76 %) in nucleotide-bound form (pdb code: 7bm8) and *parS*-bound form (pdb code: 6t1f) were used as templates for two runs. Top ranking predicted structures from these runs were subsequently used for SAXS-based structural modelling.

### Integrative structural modelling and analysis

CORAL (Petoukhov et al., 2012) from the ATSAS suite version 3.3.0-prerelease (Manalastas-Cantos et al., 2021) was used for SAXS-dependent structural modelling using averaged experimental scattering profiles. Strict two-fold symmetry was imposed for all CORAL runs. Rigid bodies of CTPBD (37-132 residues), HTHD (137-232 residues) and CTDD (239-293 residues) were used as inputs for the CORAL runs in three-domain mode. At times, combined CTPBD and HTHD (37-232 residues) and CTDD were used for CORAL runs in two-domain mode. In an attempt to improve geometry of the predicted model, positional energy minimization was performed in Phenix (Liebschner et al., 2019). Two flexible regions, that contained less than 20 % of the protein residues, were obtained from CORAL loop library. 20 CORAL runs were performed against each dataset. Additional distance restraints between the CTDD/CTPBD dimers (based on the Alphafold2 model) and between HTHD and *parS1* DNA (based on the structure of cvParBtrunc-*parS* complex, pdb code 6t1f) were added, as required.

In addition to experimental data, theoretical scattering profiles of the dimerized domains (CTDD, HTHD and CTPBD, as required) of vcParB1 were used as supplementary synthetic data during the CORAL runs. Theoretical scattering profiles were calculated from structural models of truncated dimeric domains using CRYSOL (Manalastas-Cantos et al., 2021). Segments of the predicted Alphafold2 model of vcParB1 dimer was used for calculating theoretical profiles of the dimeric CTDD and dimeric CTPBD domains. The two HTHD domains of vcParB1 were superposed on the corresponding domains of the cvParBtrunc-*parS* complex (pdb code 6t1f) for obtaining dimeric, *parS1*-bound form of HTHD for theoretical profile calculation.

The unbound form of vcParB1 and CTP-treated vcParB1 were modelled using the experimental data, and the theoretical SAXS profile of the dimeric CTDD. Two theoretical scattering curves, one for the dimeric CTDD and one for the dimeric HTHD in *parS1*-bound form, were used along with the experimental scattering data for modelling the vcParB1-*parS1* assembly. The HTHD domain of vcParB1 was altered in *Coot* (Emsley and Cowtan, 2004) to match the HTHD of cvParBtrunc-*parS* complex (pdb code 6t1f), so that it can bind *parS1* without steric conflict. The *parS1* DNA model (obtained by trimming the coordinates of the *parS* duplex, pdb code 6t1f), after shifting to the origin and reorienting the two-fold axis with the Z-axis, was kept fixed during CORAL runs. For modelling CTP-treated vcParB1 in “N-gate closed” form, theoretical SAXS profiles of dimeric CTDD and dimeric CTPBD (N-gate-closed form) were used along with the experimental scattering profile.

*Coot* (Baker et al., 2001) and Pymol (Schrodinger, LLC) were used for visualizing and superposing the structural models in graphics. APBS (Baker et al., 2001) was used for the calculation of electrostatic potential map. Amber charges and all default parameters were used for APBS calculations. Multiple sequence alignments were prepared in Jalview (Waterhouse et al., 2009) and flexible regions were predicted using the “hot loop” option in Jalview. PyMol (Schrodinger, LLC) was used for preparing figures.

### Data availability

SAXS data were deposited to SASBDB (www.sasbdb.org), with following accession numbers: SASDVD3, SASDVE3, SASDVF3, SASDVG3, SASDVH3, SASDVJ3, SASDVK3, SASDVL3, SASDVM3, SASDVN3, SASDVP3, SASDVQ3, SASDVR3, SASDVS3, SASDVT3, SASDVU3, SASDVV3.

## ACKNOWLEDGEMENTS

Department of Biotechnology (DBT), Govt. of India, provided extramural grant (BT/PR8804/BRB/10/1252/2013) and Council of Scientific and Industrial Research (CSIR), Govt. of India, provided intramural funding to support this project. BC thanks CSIR support and institutional facility. SAXS data were collected at beamline BM29 at ESRF, France, which was supported by DBT and administered by Regional Centre of Biotechnology, Faridabad. We thank BM29 staffs for help during SAXS and SEC-SAXS data collection. Anu thanks Mr. Digvijay Singh for the AUC experiment and staffs from NanoTemper technologies for DLS experiments during demonstration. Anu thanks CSIR for her fellowship. Sneh Lata thanks Indian Council of Medical Research for her fellowship.

## Supplementary Materials 1

Sources of constructs used in this study are listed in supplementary **Table S1**.

**Table S1.**
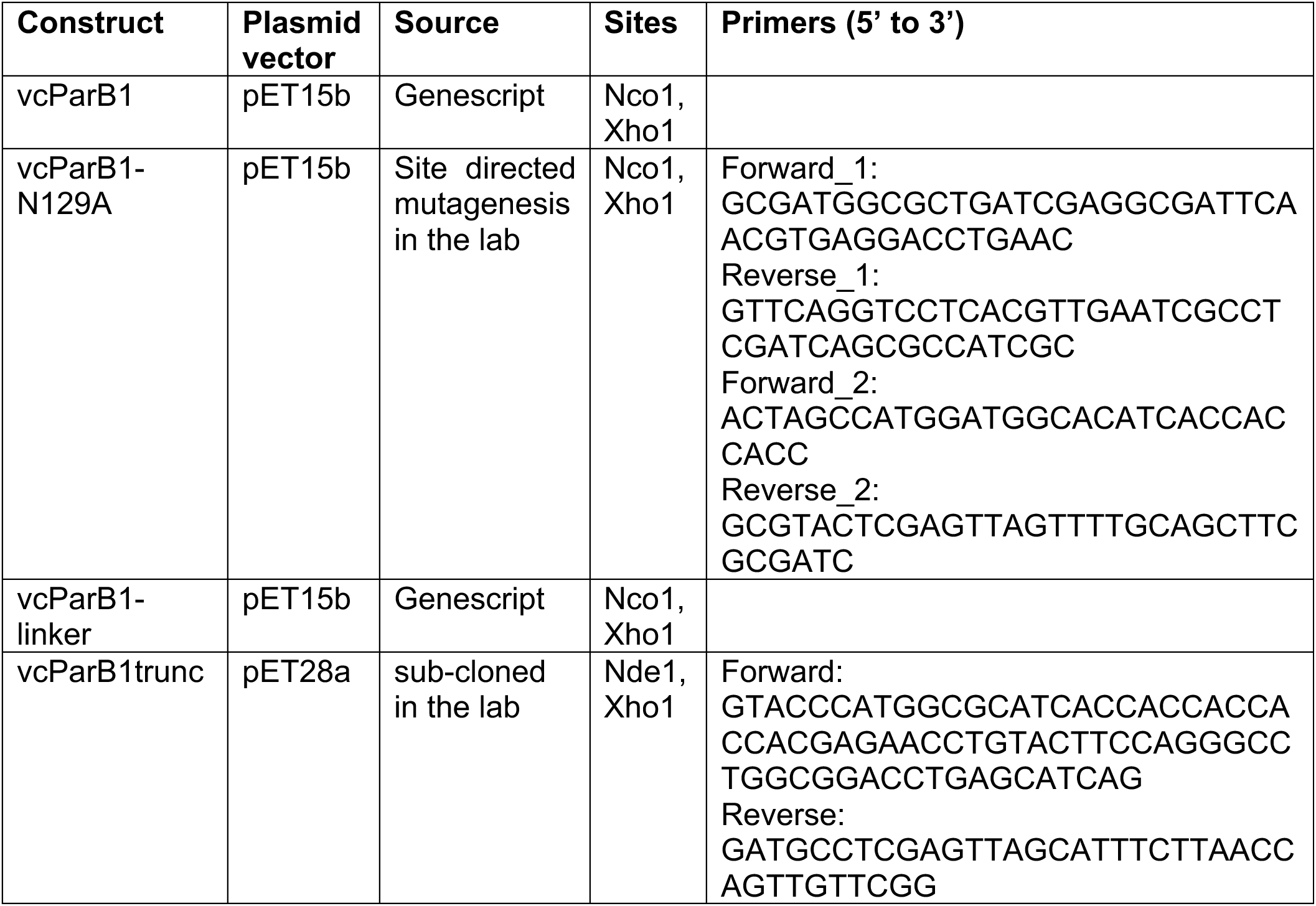
List of protein constructs.

Following materials were used for protein construct preparation: dNTP Mix (ThermoFisher scientific, cat R0192), Phusion DNA polymerase (ThermoFisher scientific, cat F530S), Fast Alkaline phosphatase (ThermoFisher scientific, cat EF0651), T4 DNA ligase (ThermoFisher scientific, cat EL0011), FastDigest Nco1 (ThermoFisher scientific, cat FD0573), FastDigest Nde1 (ThermoFisher scientific, cat FD0583), FastDigest Xho1 (ThermoFisher scientific, cat FD0694), GeneJET plasmid miniprep kit (ThermoFisher scientific, cat K0502) and GeneJET gel extraction kit, (ThermoFisher scientific, cat K0691). Polymerase chain reactions were performed in a thermocycler (Mastercycler, Eppendorf).

### Oligonucleotide sequences used in this study

<u>Oligonucleotides used for SEC-SAXS and CTPase activity assay</u>

TGTTTCACGTGAAACA (*parS1*)

GATTGCAACGATATCT (*nsDNA16*)

AGATATCGTTGCAATC (complementary *nsDNA16*)

<u>Oligonucleotides used for biolayer interferometry:</u>

Biotin-GGGATGTTTCACGTGAAACA (bt-*parS1*)

TGTTTCACGTGAAACATCCC (complementary bt-*parS1*)

Biotin-GGGAGATTGCAACGATATCT (bt-*nsDNA*)

AGATATCGTTGCAATCTCCC (complementary bt-*nsDNA*)

