## Supplemental Figure for "Combining chromatography coupled SAXS and AI-predicted structures to dissect the mechanism of ParB1-*parS1* partition assembly formation"

### Figure S1.

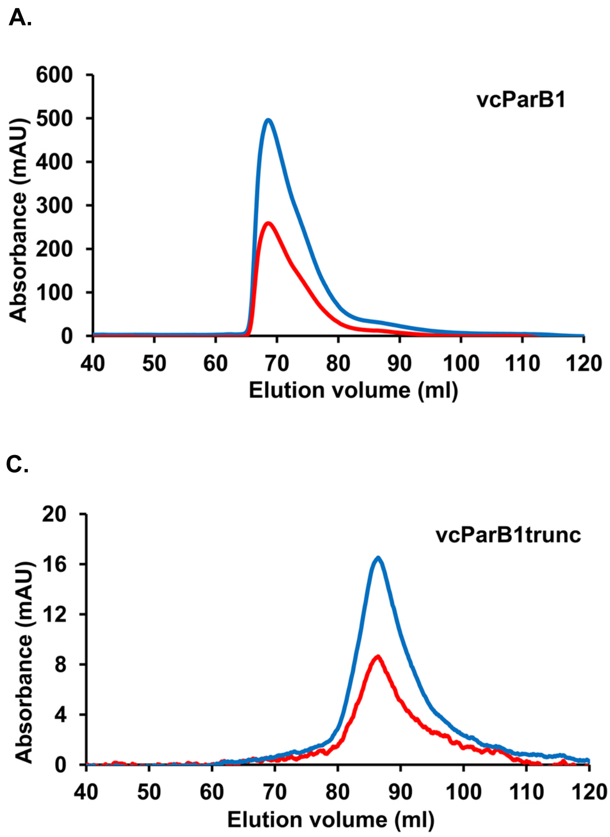


**Figure S1.** SEC elution profiles (HiLoad 16/600 superdex 200 pg column) of all the vcParB1 constructs used for SEC-SAXS studies. Absorbance at 280 nm (blue) and 260 nm (red) in mAU *versus* elution volume in ml are plotted for (A) vcParB1, (B) vcParB1-linker and (C) vcParB1trunc.

**Figure S2.**

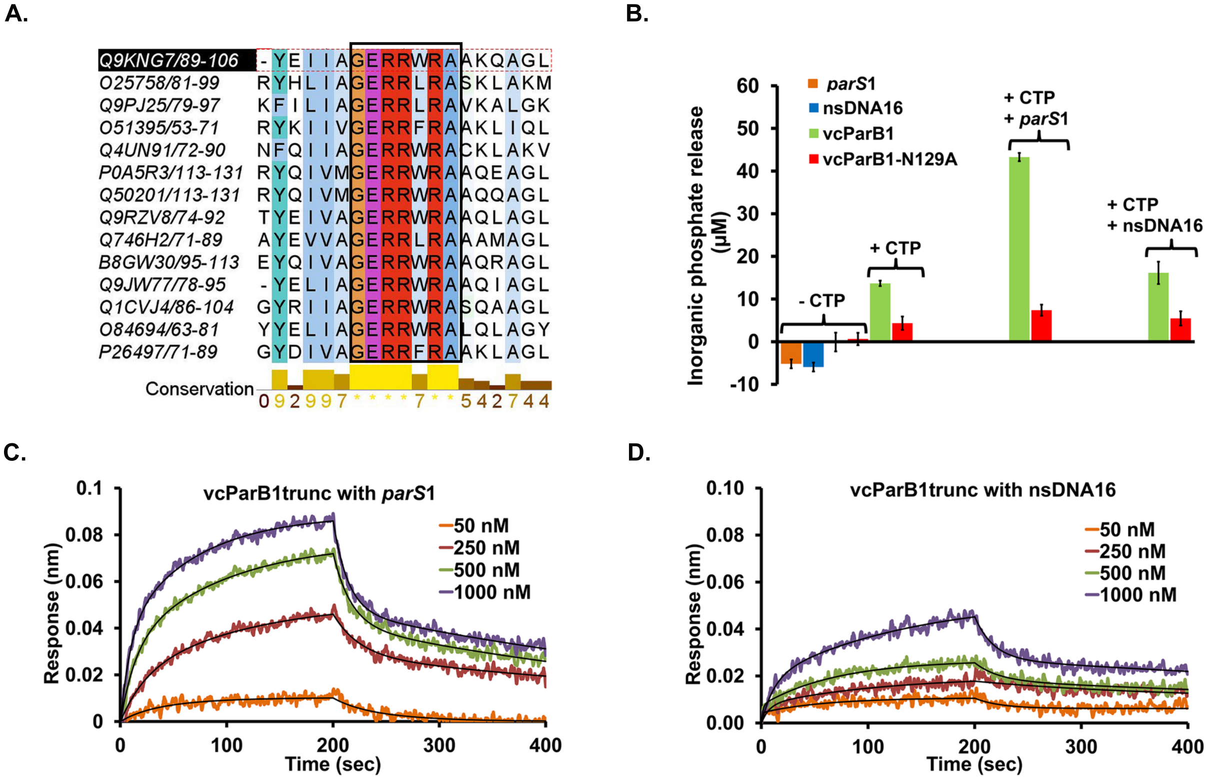
A.
Q9KNG7/89-106 -YEIIAGERRWRAAKQAGL
O25758/81-99 RYHLIAGERRLRASKLAKM
Q9PJ25/79-97 KFILIAGERRLRAVKALGK
O51395/53-71 RYKIIVGERRFRAAKLIQL
Q4UN91/72-90 NFQIIAGERRWRACKLAKV
P0A5R3/113-131 RYQIVMGERRWRAAQEAGL
Q50201/113-131 RYQIVMGERRWRAAQQAGL
Q9RZV8/74-92 TYEIVAGERRWRAAQLAGL
Q746H2/71-89 AYEVVAGERRLRAAAMAGL
B8GW30/95-113 EYQIVAGERRWRAAQRAGL
Q9JW77/78-95 -YELIAGERRWRAAQIAGL
Q1CVJ4/86-104 GYRIIAGERRWRASQAAGL
O84694/63-81 YYELIAGERRWRALQLAGY
P26497/71-89 GYDIVAGERRFRAAKLAGL
Conservation
0 9 2 9 9 7 * * * * 7 * * 5 4 2 7 4 4
B.
Inorganic phosphate release (µM)
60
50
40
30
20
10
0
-10
parS1
nsDNA16
vcParB1
vcParB1-N129A
- CTP
+ CTP
+ CTP
+ parS1
+ CTP
+ nsDNA16
C.
vcParB1trunc with parS1
Response (nm)
0.1
0.08
0.06
0.04
0.02
0
0
100
200
300
400
Time (sec)
50 nM
250 nM
500 nM
1000 nM
D.
vcParB1trunc with nsDNA16
Response (nm)
0.10
0.08
0.06
0.04
0.02
0.00
0
100
200
300
400
Time (sec)
50 nM
250 nM
500 nM
1000 nM

**Figure S2.** CTPase activity and DNA-binding of vcParB1. (A) Multiple sequence alignment of an N-terminal region of vcParB1 (Uniprot accession number Q9KNG7, residues 89 to 106, encompassing the conserved GxxRxxA motif present in ParB CTPases, 3) with homologous ParBs (*Helicobacter pylori*: O25758, *Campylobacter jejuni*: Q9PJ25, *Borreliella burgdorferi*: O51395, *Rickettsia felis*: Q4UN91, *Mycobacterium bovis*: P0A5R3, *Mycobacterium leprae*: Q50201, *Deinococcus radiodurans*: Q9RZV8, *Thermus thermophilius*: Q746H2, *Caulobacter vibrioides*: B8GW30, *Neisseria meningitides*: Q9JW77, *Myxococcus xanthus*: Q1CVJ4, *Chlamydia trachoatis*: O84694, *Bacillus subtilis*: P26497) is shown. (B) Bar graph showing the results of CTPase activity assay of vcParB1 (green, dimer concentration 4 μM) and vcParB1-N129A mutant (red, dimer concentration 4 μM), with and without added 16-meric specific, palindromic DNA duplex (*parS1,* 4 μM) and 16-meric non-specific DNA duplex (nsDNA16, 4 μM). Results of the control experiments without CTP are shown alongside. (C-D) Sensograms (response in nm *versus* time in sec) of interactions of a truncated version of vcParB1 containing the CTPBD and HTHD domains (vcParB1trunc) with (C) *parS1,* and with (D) nsDNA16 as a control, are shown with fitted curves (black solid line). DNA sequences used in the CTPase activity assay and bio-layer interferometry experiments are described in supplementary material S1.

**Figure S3.**

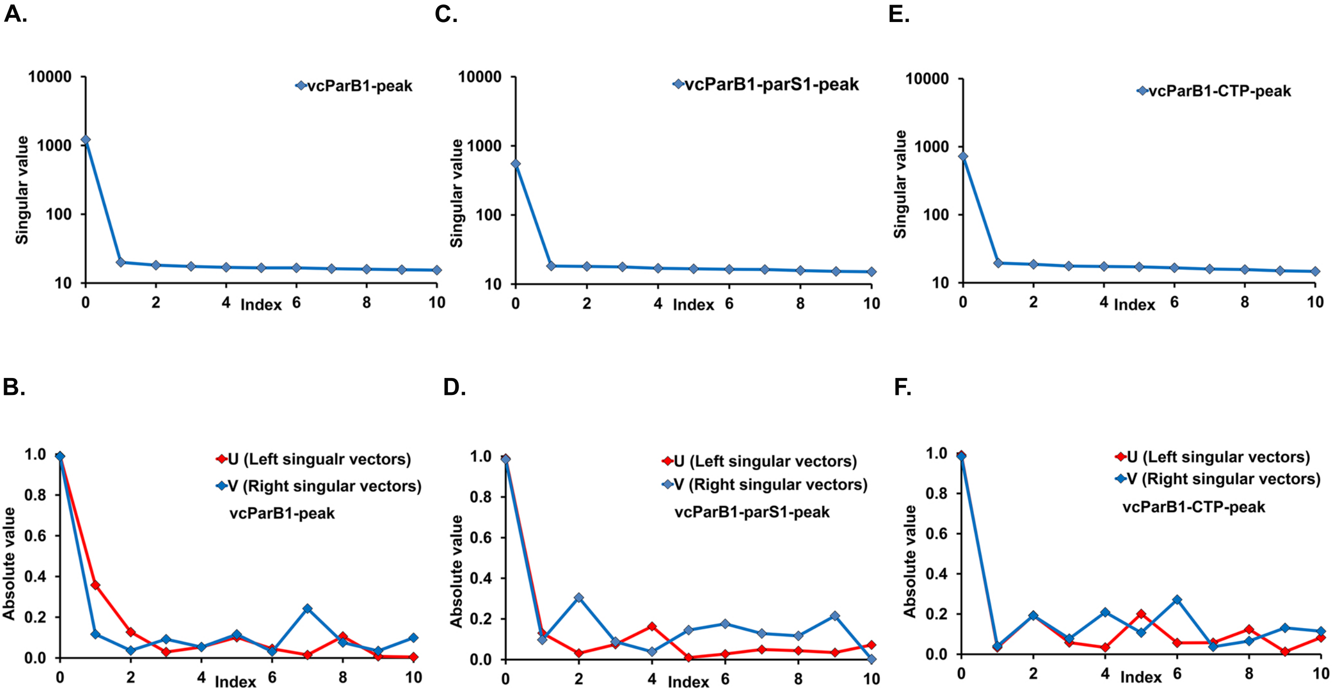


**Figure S3.** Singular value decompositions of the SEC-SAXS elution profiles of (A-B) vcParB1, (C-D) vcParB1-*parS1*, (E-F) vcParB1-CTP peaks. Plots of singular values *versus* index are shown in the top panel and autocorrelation plots of left (red) and right (blue) singular vectors *versus* index are shown in the bottom panel for all profiles (computed in BioXtas RAW, Hopkins, 2024). The SVD analyses were performed on the frames that were used for averaging to obtain the peak profiles for structural modelling.

### Figure S4.

A.

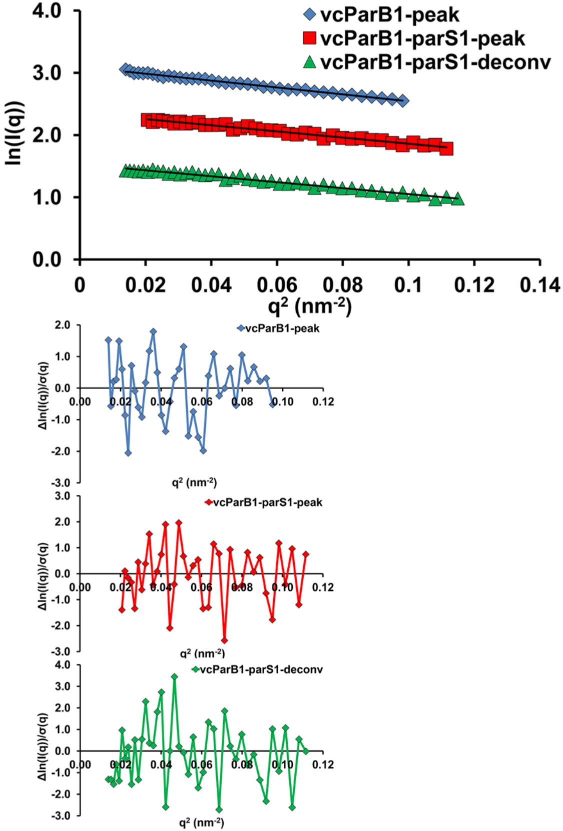


B.

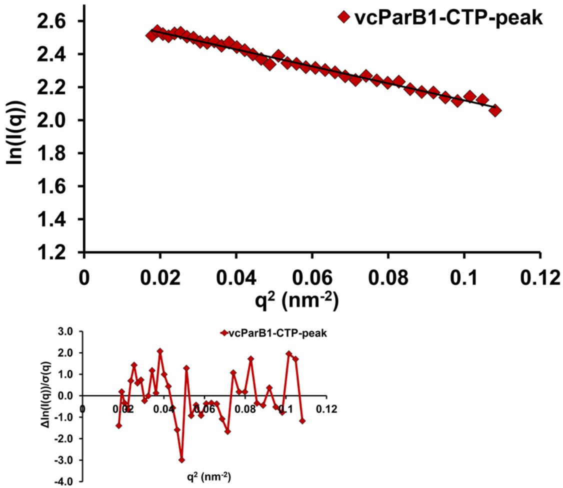


**Figure S4.** Guinier plots (lnI(q) *versus* $q^2$, where I(q) is intensity in arbitrary unit and q is momentum transfer in $nm^{-1}$, $qR_g \leq 1.3$), and corresponding residuals, are shown for (A) vcParB1-peak (blue diamond), vcParB1-*parS1*-peak (red square), vcParB1-*parS1*-deconv (green triangle) and (B) vcParB1-CTP-peak (brown diamond) profiles.

#### Figure S5.

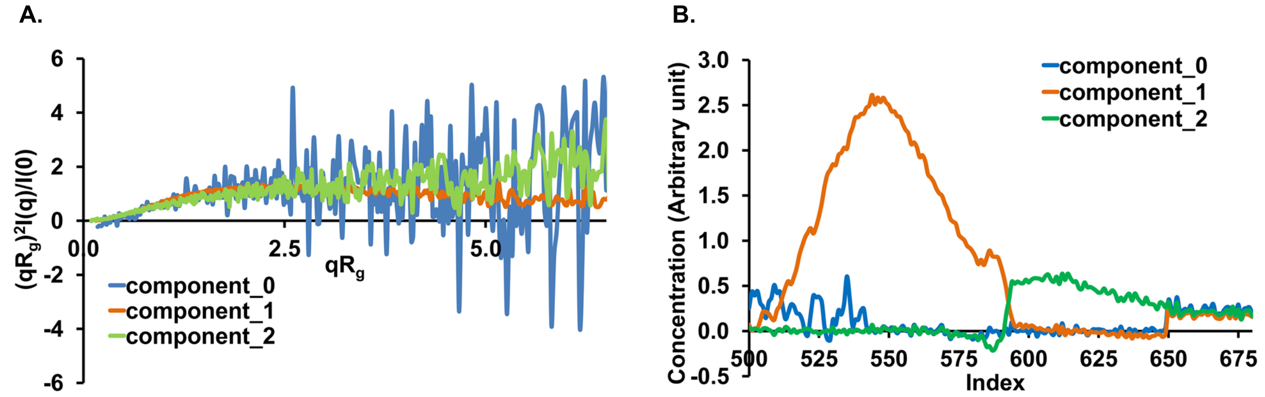


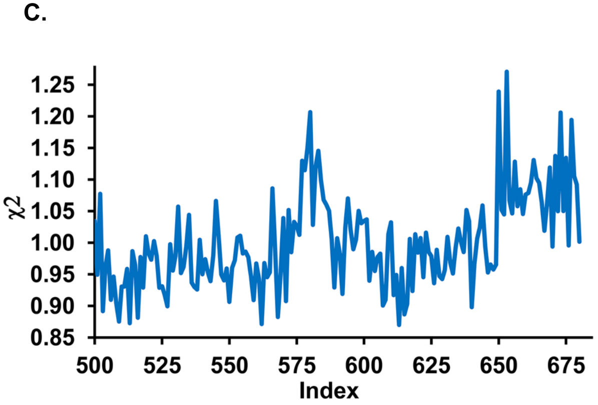


**Figure S5.** REGALS (Meisburger et al., 2021) decomposition of vcParB1-*parS1* SEC-SAXS profile. (A) Dimensionless Kratky plot $(I(q)/I(0).(qR_g)^2$ *versus* $qR_g)$ of the REGALS deconvoluted profiles (3 components in blue, orange and green) of vcParB1-*parS1* SEC-SAXS data surrounding the peak region. (B) Concentration (in arbitrary unit) plots of the three components (in blue, orange and green) obtained from REGALS analysis. (C) Mean error-weighted $\chi^2$ *values* (blue) *versus* index numbers from REGALS analysis. The component 1 (vcParB1-*parS1*-deconv, orange) was used for subsequent analysis.

**Figure S6.**

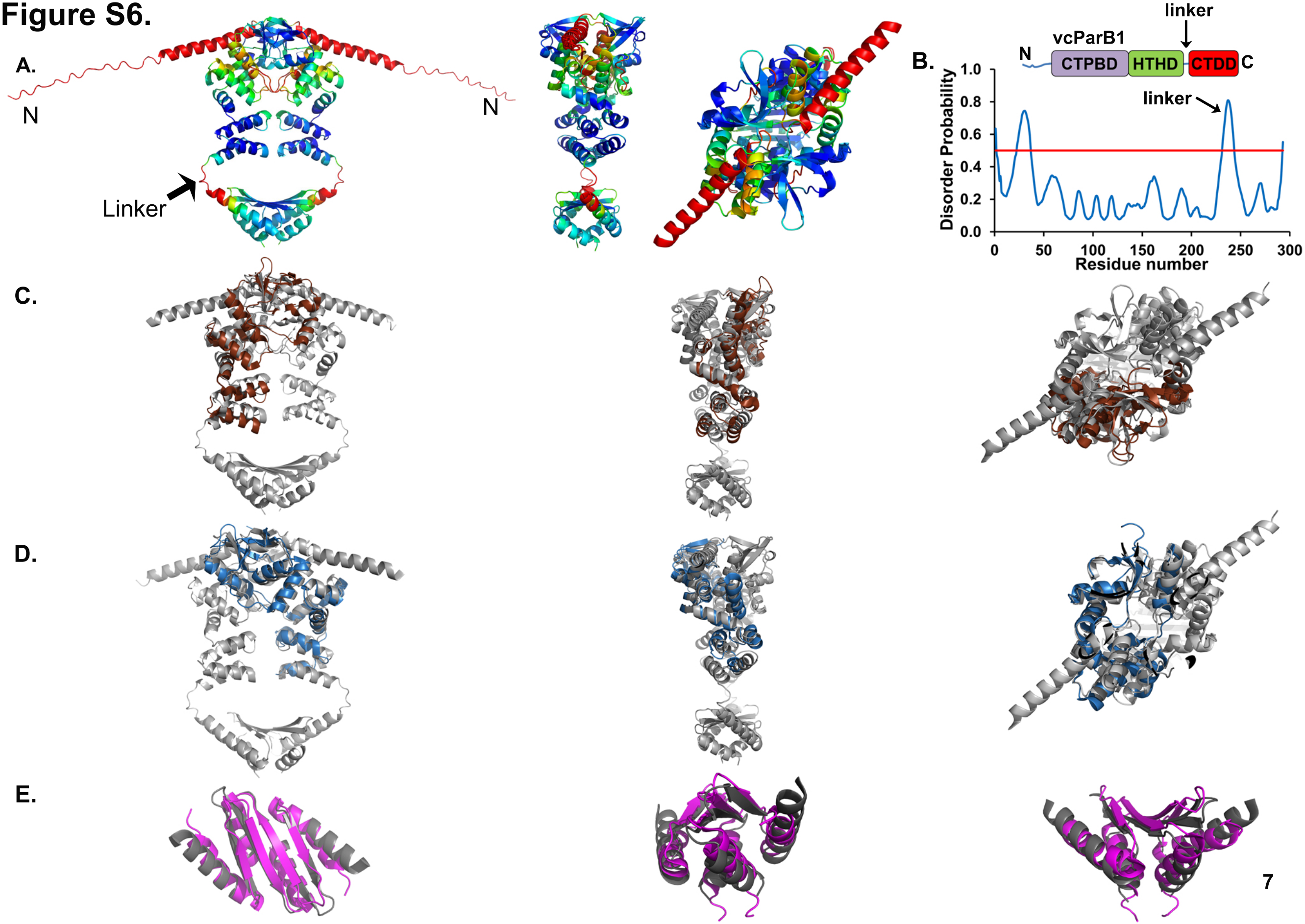

A.
N
N
Linker
B.
vcParB1
linker
N
CTPBD
HTHD
CTDD
C
linker
Disorder Probability
1.0
0.8
0.6
0.4
0.2
0.0
0
50
100
150
200
250
300
Residue number
C.
D.
E.

**Figure S6.**

F.

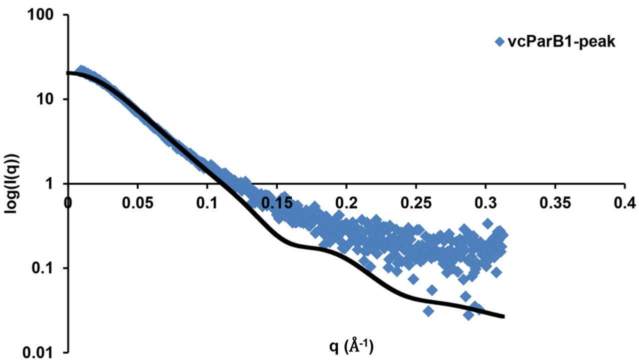


G.

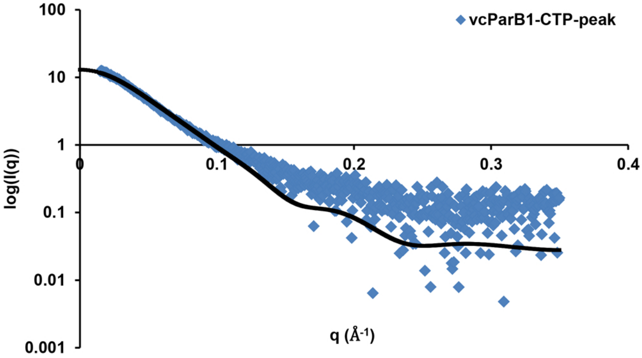


H.

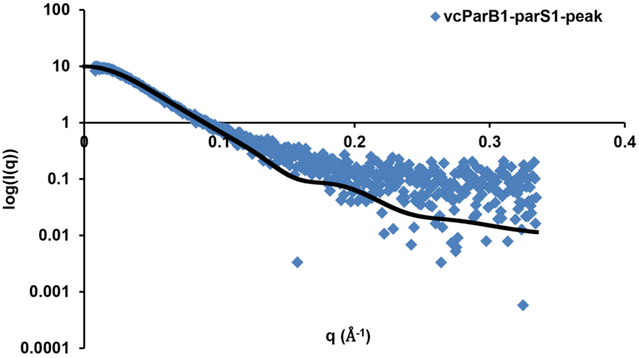

**Figure S6.** Alphafold multimer predicted structures of dimeric, full-length vcParB1. (A) Three views of predicted structure of vcParB1 colored according to the pLDDT score (90 to 50, blue to red) in cartoon. The N-terminal disordered region is shown only in the first view for clarity. (B) Schematic diagram in the upper panel shows the domain organization of full-length vcParB1. Predicted intrinsic disorder (disorder probability *versus* residue number, calculated in PrDOS, prdos.hgc.jp) of vcParB1 plotted against residue numbers is shown in the lower panel. The threshold is shown as a horizontal red line. (C-D) Three different views of the predicted structures of vcParB1 (grey; the N-terminal disordered region is not shown) superposed with (C) *parS1*-bound cvParBtrunc (brown, pdb code: 6t1f, RMSD 3.4 Å for 182 Cα atoms aligned), (D) a nucleotide-bound form of cvParBtrunc (light blue, pdb code: 7bm8, RMSD 2.2 Å for 171 Cα atoms aligned). (E) The dimeric CTDD region of predicted vcParB1 (grey cartoon) aligned with a model of the C-terminal domain of bsParB (magenta cartoon, pdb code: 5noc, RMSD 1.5 Å for 48 Cα atoms aligned). All figures show Alphafold2 structures of vcParB1 predicted using the nucleotide- bound form of cvParBtrunc (pdb code: 7bm8) as a custom template, except S6C. In S6C, the Alphafold2 structure of vcParB1 that was predicted using *parS*1-bound cvParBtrunc (pdb code: 6t1f) as a custom template is shown. (F-H) Theoretical scattering profile (solid black line) of the Alphafold2 structure of vcParB1 used for modelling is shown (logI(q) *versus* q in $Å^{-1}$), fitted to experimental (F) vcParB1-peak ($\chi^2$ ~ 9.8), (G) vcParB1-CTP-peak ($\chi^2$ ~ 4.7) (H) vcParB1-*parS1*-peak ($\chi^2$ ~ 2.5) scattering profiles (blue squares). Note that the predicted Alphafold2 model of vcParB1 does not include *parS1*.

### Figure S7.

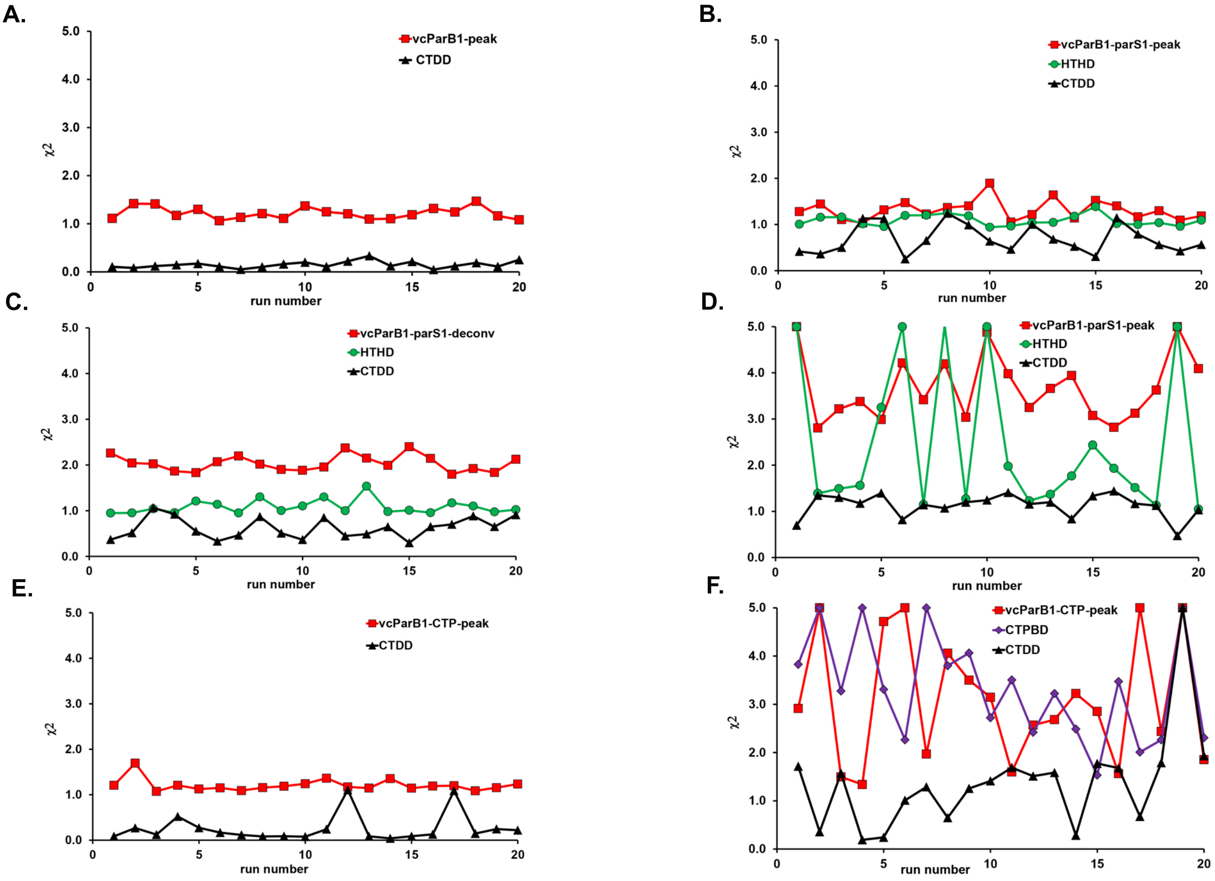

A.
vcParB1-peak
CTDD
χ2
run number
B.
vcParB1-parS1-peak
HTHD
CTDD
χ2
run number
C.
vcParB1-parS1-deconv
HTHD
CTDD
χ2
run number
D.
vcParB1-parS1-peak
HTHD
CTDD
χ2
run number
E.
vcParB1-CTP-peak
CTDD
χ2
run number
F.
vcParB1-CTP-peak
CTPBD
CTDD
χ2
run number

**Figure S7.** The $\chi^2$ values of the 20 CORAL runs against experimental and synthetic SAXS profiles. Plots of $\chi^2$ values against run numbers for hybrid modelling of vcParB1, vcParB1-*parS1* and vcParB1-CTP, are shown as follows: (A) vcParB1 modelled against vcParB1-peak data (red) and synthetic data obtained from the CTDD dimer (black); (B) vcParB1-*parS1* modelled against vcParB1-*parS1*-peak data (red), synthetic data obtained from the CTDD dimer (black), and synthetic data obtained from HTHD dimer in *parS*-bound form (green); (C) vcParB1-*parS1* modelled against vcParB1-*parS1*-deconv data (red), synthetic data obtained from the CTDD dimer (black), and synthetic data obtained from HTHD dimer in *parS*-bound form (green); (D) vcParB1-*parS1* modelled against vcParB1-*parS1*-peak data (red), synthetic data obtained from the CTDD dimer (black), synthetic data obtained from the HTHD dimer in *parS*-bound form (green), and additional distance restraints for positioning the CTPBD domains closer; (E) CTP-bound vcParB1 modelled against vcParB1-CTP-peak data (red) and synthetic data obtained from the CTDD dimer (black); (F) CTP-bound vcParB1 modelled against vcParB1-CTP-peak data (red), synthetic data obtained from the CTDD dimer (black), and CTPBD dimer in “N-gate closed” form (violet). All $\chi^2$ values above 5 were set equal to 5 for visualization purpose.

### Figure S8.

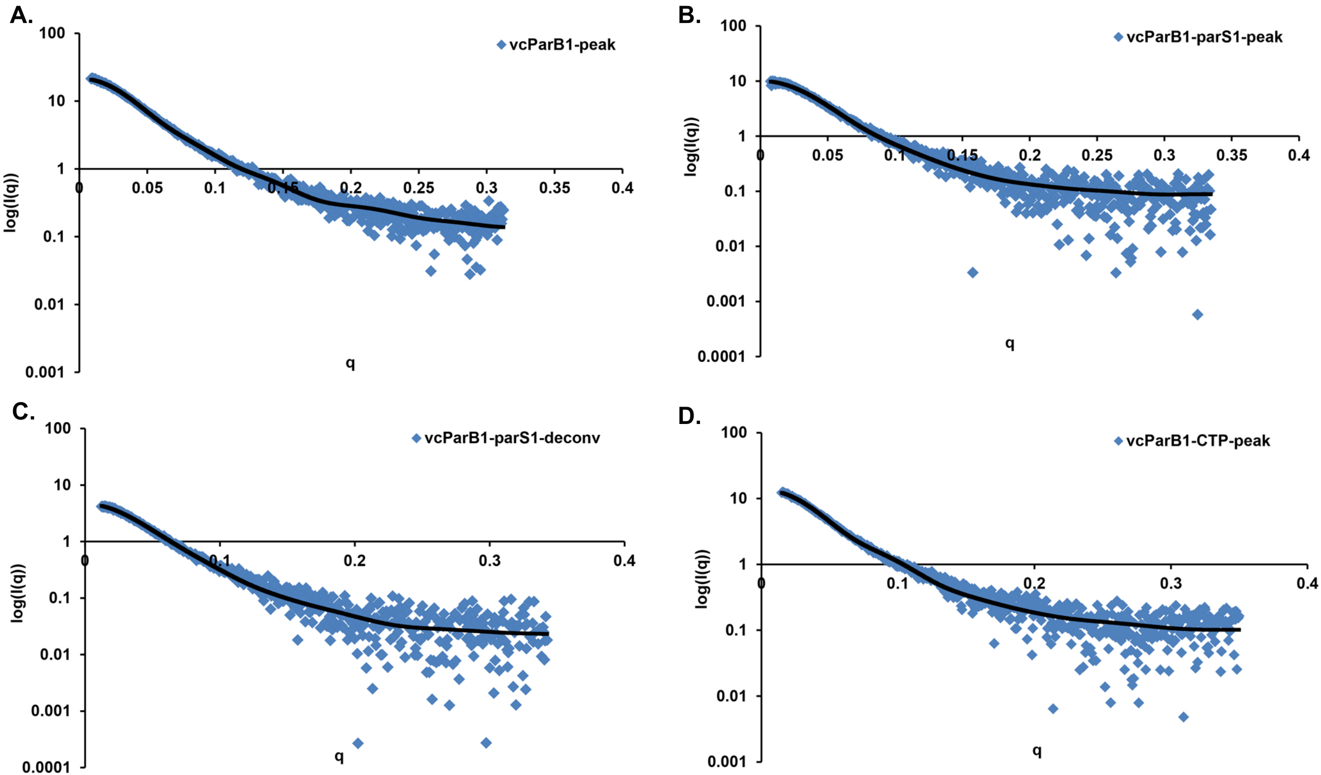


**Figure S8.** Plots of logI(q) *versus* q (Å$^{-1}$) showing fits between experimental profiles (blue diamonds) and theoretical profiles calculated from representative hybrid models (black line) for (A) vcParB1-peak ($\chi^2$ ~1.4), (B) vcParB1-*parS1*-peak ($\chi^2$ ~1.2), (C) vcParB1-*parS1*-deconv ($\chi^2$ ~2.0) and (D) vcParB1-CTP-peak ($\chi^2$ ~1.3) datasets.

#### Figure S9.

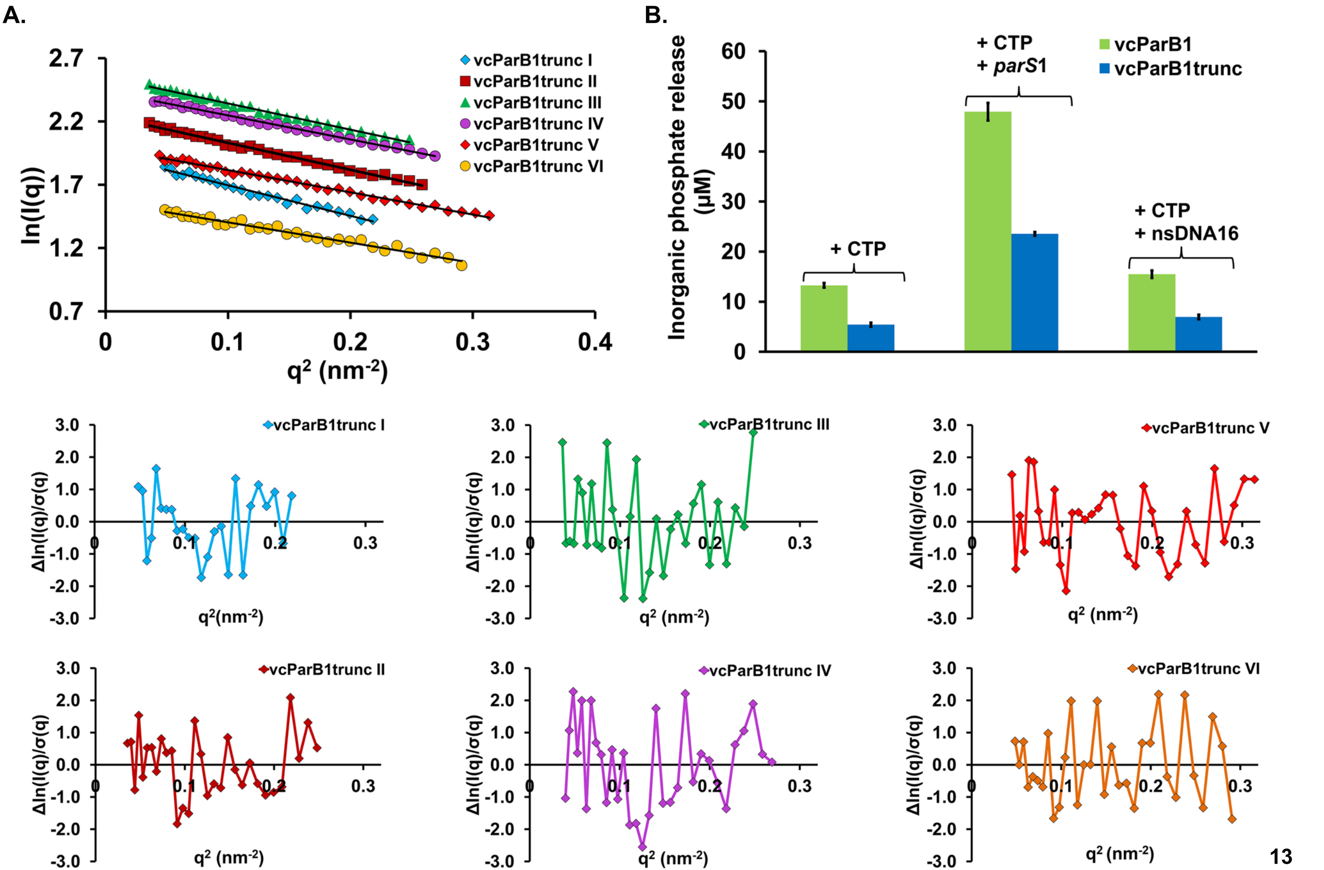
A.
B.
ln(I(q))
q² (nm⁻²)
vcParB1trunc I
vcParB1trunc II
vcParB1trunc III
vcParB1trunc IV
vcParB1trunc V
vcParB1trunc VI
Inorganic phosphate release (µM)
vcParB1
vcParB1trunc
+ CTP
+ CTP + parS1
+ CTP + nsDNA16
Δln(I(q)/σ(q))
q²(nm⁻²)

**Figure S9.** (A) Guinier plots (lnI(q) *versus* $q^2$, $R_g \leq 1.3$) of the segments of SAXS profiles obtained from the SEC-SAXS datasets of vcParB1trunc, with the corresponding residuals (lower panel), are shown. (B) Bar graph showing the CTPase activity of vcParB1 (green), and vcParB1trunc (blue), in the presence and absence of specific and non specific DNA (protein dimer concentration: 4 μM, duplex DNA concentration: 1 μM).

### Figure S10.

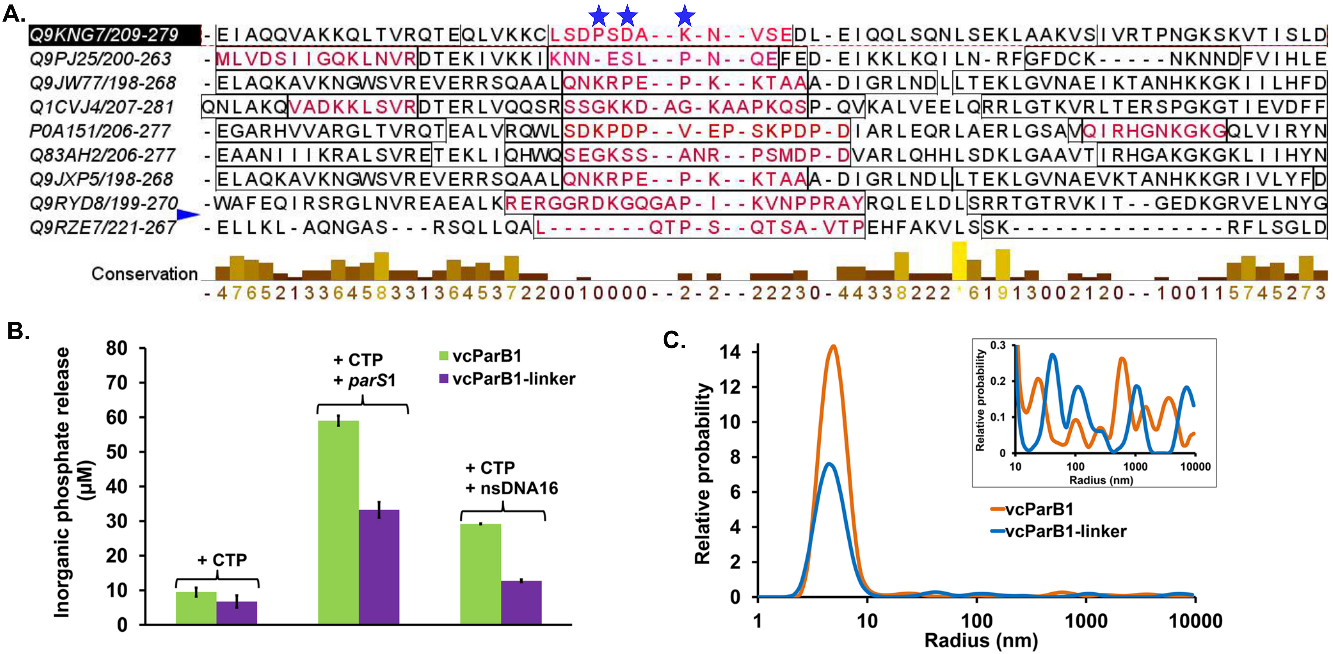


**Figure S10.** (A) Multiple sequence alignment of the linker region of vcParB1 (Uniprot accession number: Q9KNG7, residues 209-279) and other ParB homologs (Uniprot accession numbers: Q9PJ25, Q9JW77, Q1CVJ4, P0A151, Q83AH2, Q9JXP5, Q9RYD8, Q9RZE7) is shown with predicted flexibility (hot loop, coloured in red). Sequence conservation scores are shown below the sequence alignment. The positions of the three mutations in the vcParB1-linker construct are marked with blue stars. (B) Bar graph showing the CTPase activity of vcParB1 (green) and vcParB1-linker (violet), with and without specific and non specific DNA (protein dimer concentration: 4 μM, duplex DNA concentration: 4 μM). (C) Plot of relative probability *versus* hydrodynamic radius (nm) for vcParB1 (orange, cumulative polydispersity index 0.14) and vcParB1-linker (blue, cumulative polydispersity index 0.38), at 1 mg/ml (~29 μM) concentrations. Inset is showing a magnified version of this plot in the hydrodynamic radius range of 10-10000 nm.
